# Distinct structure and gating mechanism in diverse NMDA receptors with GluN2C and GluN2D subunits

**DOI:** 10.1101/2022.11.09.514853

**Authors:** Jilin Zhang, Ming Zhang, Qinrui Wang, Han Wen, Zheyi Liu, Fangjun Wang, Yuhang Wang, Fenyong Yao, Nan Song, Zengwei Kou, Yang Li, Fei Guo, Shujia Zhu

**Author notes:** Correspondence: Dr. Shujia Zhu. These authors contributed equally (Jilin Zhang, Ming Zhang, Qinrui Wang, Han Wen).

## Abstract

*N*-Methyl-D-Aspartate (NMDA) receptors are essential for many brain functions. These receptors are heterotetramers typically comprising two GluN1 subunits and two GluN2 subunits. The latter could alternate among four subtypes (N2A-N2D) and determine the functional diversity of NMDA receptors^1, 2^. For example, receptors containing N2C or N2D exhibit 50-fold lower channel open probability (Po) than those containing N2A (ref.^3–5^). Structures of N2A- and N2B-containing receptors have been extensively characterized, providing molecular basis for understanding NMDA receptor function^6–14^. Here we report the cryo-EM structures of N1-N2D and N1-N2C di-heterotetramers (di-receptors), and N1-N2A-N2C tri-heterotetramer (tri-receptor) at a resolution up to 3.0 Å. Structural analysis showed that the bilobate N-terminal domain (NTD) in N2D adopted an intrinsic closed conformation, leading to a compact NTD tetramer in N1-N2D receptor. Functional studies further demonstrated that, in di-receptors containing N2D but not N2A or N2B, crosslinking NTD at the tetrameric interface had no effect on channel activity, while crosslinking ligand-binding domain (LBD) of two N1 protomers significantly elevated Po. Surprisingly, we found that the N1-N2C di-receptors spontaneously oscillated between symmetric and asymmetric conformation. The later one occupied a predominant population, whereby two N2C protomers exhibited distinct conformation. This asymmetry, which was also found to a lesser extent in N1-N2A di-receptor^10^, was further locked by the binding of an N2C-specific allosteric potentiator PYD-106 to a unique binding pocket between NTD and LBD in only one N2C protomer. Finally, N2A and N2C in the N1-N2A-N2C tri-receptor displayed the conformation close to that found in one protomer of N1-N2A and N1-N2C di-receptors, respectively. These findings provide a comprehensive structural understanding of diverse functional properties of major NMDA receptor subtypes.

## Main

NMDA receptors display functional heterogeneity with N2C and N2D subtypes significantly distinct from N2A and N2B subtypes in their gating activity, biophysical and pharmacological properties^2, 15, 16^. The N2C and N2D subunits show close evolutionary conservation^17^, with N2C- and N2D-containing receptors exhibiting low channel activity, high agonist potency, less magnesium block, and reduced calcium permeability, as compared to those of N2A- and N2B-containing receptors^2^. While N2A and N2B are dominantly expressed in the cerebral cortex and hippocampus, N2D is more restricted to the thalamus and hypothalamus, and N2C mainly exists in the cerebellum and olfactory bulb^18^. At the cellular level, N2C is predominately expressed in cerebellar granule cells^18, 19^, and both N2C and N2D are enriched in the GABAergic interneurons^20, 21^. Under physiological conditions, both N2C- and N2D-containing NMDA receptors are crucial for excitation-inhibition balance of neuronal activity^22^. Dysfunctions of these receptors are involved in neurological and psychiatric diseases^23–28^. For instance, astrocytic N2C-containing NMDA receptors in nucleus accumbens mediate cocaine preference and neuroadaptations^29^, N2D-containing receptors influence emotional behaviors through the regulation of cell-specific synaptic transmission^30^. Therefore, molecules selectively targeting to the N2C- and N2D-containing NMDA receptors could be powerful tools for probing brain functions involving these receptor subtypes with regional and cell-type specificity. To date, the full-length structures and gating mechanisms of N2C- and N2D-containing NMDA receptors remain elusive. By combining cryo-EM, *in silico* calculation, mass spectrometry, biochemistry and single-channel recording, we here explored structural architecture, gating transition and subtype-specific pharmacology in N1-N2D and N1-N2C di-receptors and N1-N2A-N2C tri-receptor.

### Intact N1-N2D receptor structures and functional transition

First, we co-expressed constructs encoding the C-terminal domain (CTD) truncated human N2D and N1a (without RNA splicing exon 5) in HEK293S cells, and purified tetrameric N1a-N2D di-receptors (Extended Data Fig.1a-d). We firstly solved the cryo-EM structures of N1a-N2D receptor in complex with co-agonists glycine and glutamate (Gly-Glu), or with glycine and competitive antagonist R-CPP (Gly-CPP) (Fig.1a, b). During data processing, application of C1 or C2 symmetry for 3D refinement yielded two maps with similar conformation (Extended Data Fig.1e, f). We thus speculated that N1-N2D di-receptor is likely to adopt an overall two-fold symmetry, perpendicularly to the plane of plasma membrane. Our final cryo-EM maps with C2 symmetry yielded at 4.0 Å and 3.7 Å resolution for Gly-Glu and Gly-CPP states, respectively (Extended Data Fig.1g, h).

Globally similar to the bouquet-shaped N1-N2A and N1-N2B di-receptors^6–11^, N1-N2D receptor is assembled as a heterotetramer with two N1 peripheral and two N2D proximal to the central axis, top-down viewed from NTD layer (Fig.1a, b). Topologically, the clamshell-liked NTD (R1 plus R2 lobes) was distal to the membrane on the top, the transmembrane domain (TMD) embedded in the lipid bilayer, and sandwiched in-between the bi-lobe LBD (D1 plus D2) harboring the agonist binding pocket. The swapping feature of dimer association between NTDs and LBDs^6, 7^ was also present in N1-N2D receptors. In both maps, we could visualize the electron densities at the cleft of N1-LBD for Gly, and of N2D-LBD for Glu and R-CPP (Fig.1c). Both maps enabled us to build most of the secondary structures, including TM3 and TM2 helices that formed the ion channel gate and selective filter (Fig.1d). HOLE analysis revealed that ion channel gate in Gly-Glu state and Gly-CPP state with a smallest radius of ∼2 Å and less than 1 Å (Fig. 1e), indicating that the Gly-Glu and Gly-CPP bound receptors were likely to be trapped in the pre-open and inhibited states, respectively. We also determined the structure of N1b-N2D receptor in Gly-Glu bound state with the presence of N1 splicing cassette exon 5 at 5.1 Å resolution (Fig.1f and Extended Data Fig.2). Electron density of exon 5 cassettes was visible between α-helix 6 and β-sheet 7 on N1-NTD, similar to that found in N1b-N2B receptor structure. Compared with N1a-N2D receptor, the exon 5 polypeptide enlarged the edge distances of the triangle comprised by N1-R2, N1-D1 and N2D-D1 (Fig.1g).

**Fig. 1.**
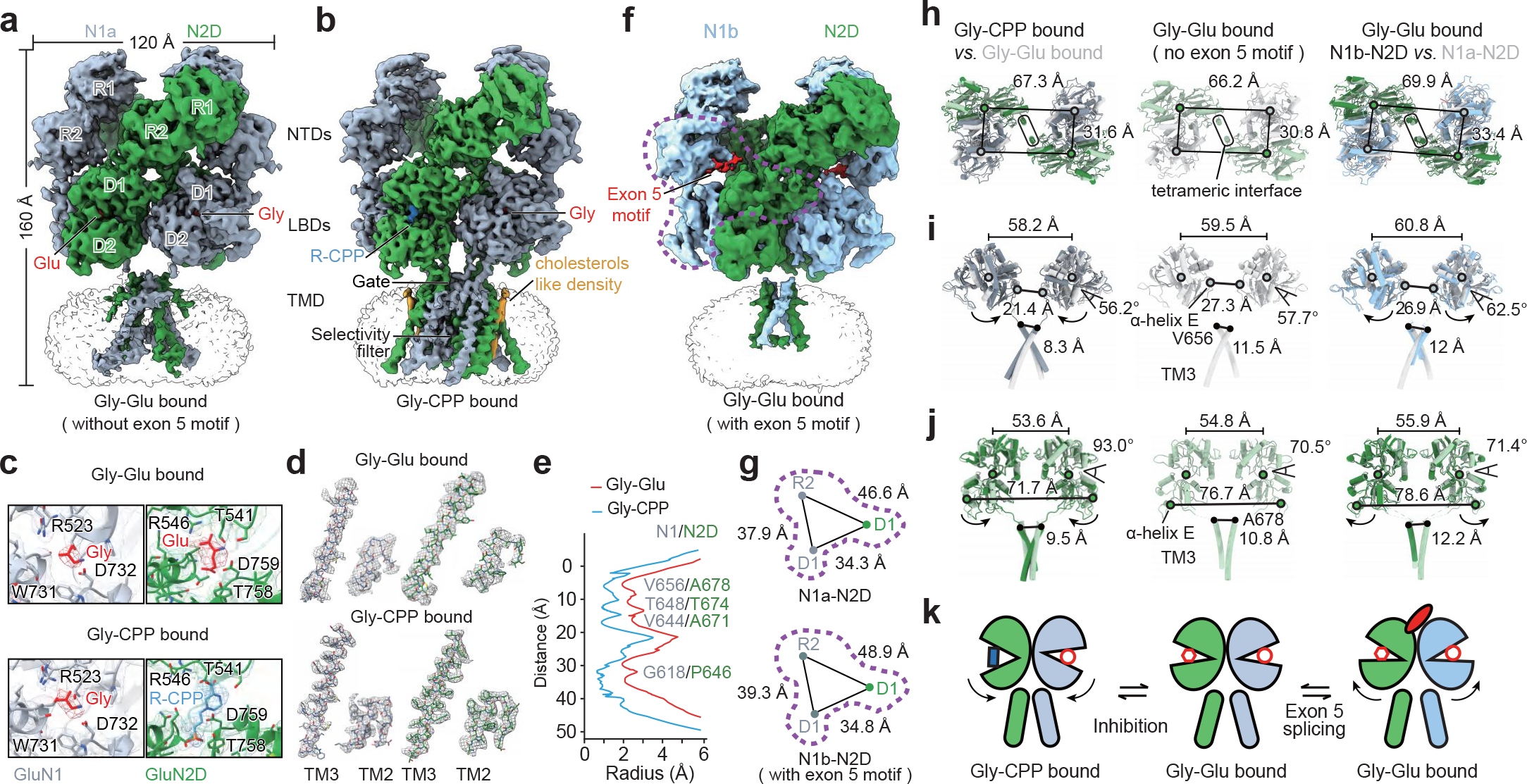
Molecular architecture and functional transition of N1-N2D receptor. **a-b**, Cryo-EM structures of N1-N2D receptor in Gly-Glu bound (**a**) and Gly-CPP bound (**b**) state. Electron density of agonists glycine and glutamate (abbreviated as Gly and Glu, in red) and antagonist R-CPP (in blue) are highlighted. **c,** Zoom-in views of N1a- (in gray) and N2D-LBD (in green) cleft with EM density and structural coordinate for Gly (in red), Glu (in red), R-CPP (in cyan) and key binding residues. **d**, Electron densities of TM2 and TM3 helices for N1a and N2D of N1a-N2D receptors in Gly-Glu and Gly-CPP states. **e**, HOLE analysis of the N1a-N2D receptor in Gly-Glu, Gly-CPP states. Pore residuals of N1a and N2D are represented next to the corresponding location. **f**, Cryo-EM structure of N1b-N2D receptor in Gly-Glu bound state, with the presence of exon 5 motif (with polypeptide of D205-P210 visible) colored in red. The exon 5 motif binding interface formed by N1-R2, N1-D1 and N2D-D1 lobes were highlighted by violet circle. **g**, Diagram illustration for the center-of-mass (COM) distance of triangle geometry connecting the N1-R2, N1-D1 and N2D-D1 lobes, with (lower panel) or without (upper panel) exon 5 motif. **h-j**, Structural analysis of top-down viewed tetrameric NTD (**h**), of side-viewed two N1 (**i**) and N2D (**j**) protomers. Center-of-mass (COM) of each lobe, domain and α-helix E (P670-R673 for N1 and R696-Q699 for N2D) is shown in empty circle. Cα atoms of N1^A652^ and N2^A678^ in gate are marked in filled circle. Rounded rectangles in (**h**) indicate the NTD tetrameric interface. The dihedral angles for indicating opening-closure degree of LBD are assessed by connecting the Cα of I403, S688, V735, A715 in N1, and P124, E525, S309, E169 in N2D, respectively. Arrows indicate the conformational changes of Gly-CPP or the exon 5 motif modulated state receptors compared to the Gly-Glu state. **k,** Cartoon illustration for the conformational transition of N1-N2D receptor among Gly-Glu, Gly-CPP and the exon 5 motif modulated states.

**Fig. 2.**
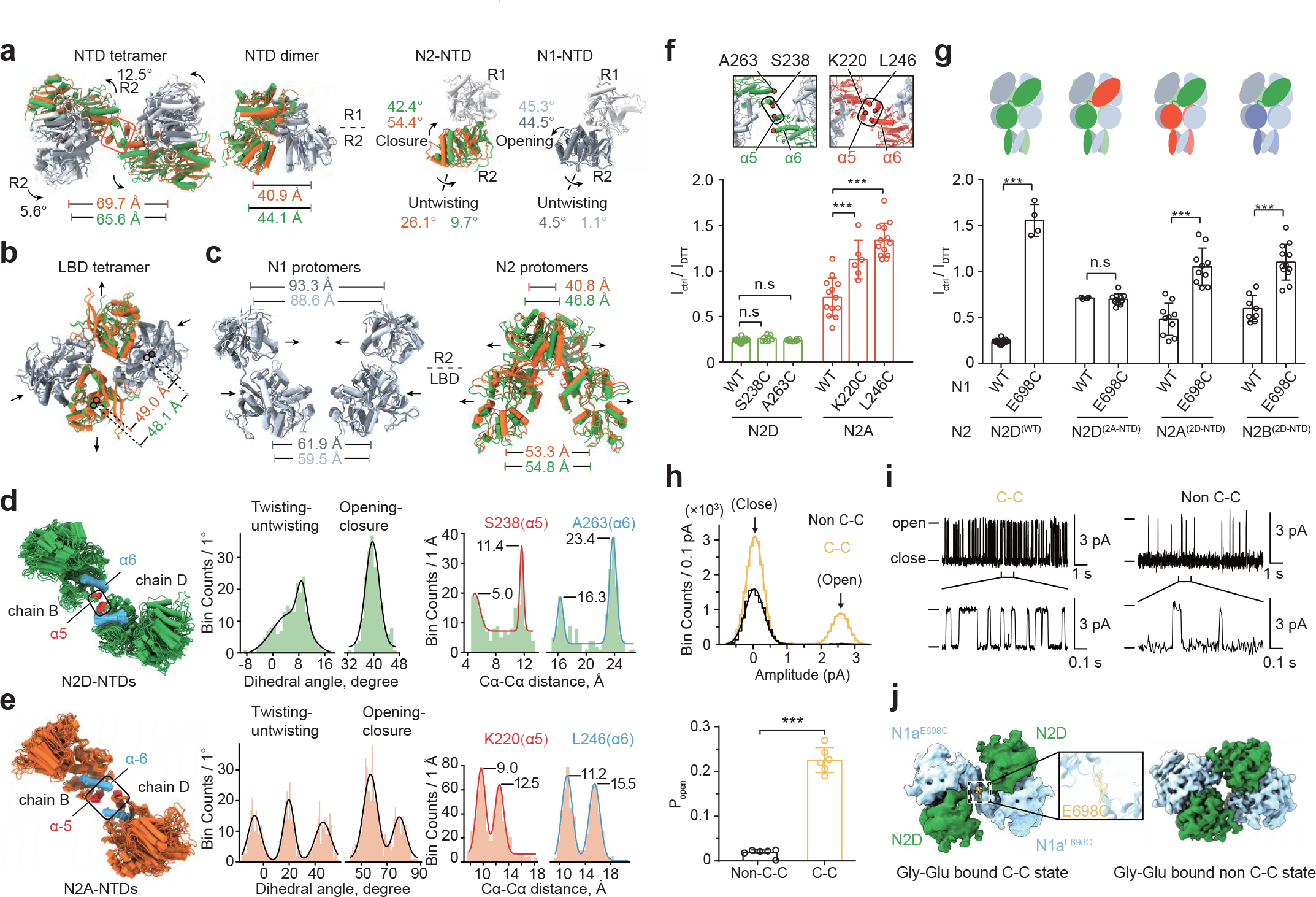
NTD and LBD function in N1-N2D receptor. **a-c**, Conformation comparison of NTD tetramer and heterodimer (left two), N1- and N2-NTD protomers (superimposed with R1 lobes, right two), LBD tetramer (**b**) and two N1 and N2 protomers (**c**) between Gly-Glu bound N1-N2D and N1-N2A (PDB:6MMP, ref.^10^) di-receptors. Rounded rectangles in (**a**) indicate the tetrameric interface of N1-N2A (orange) and N1-N2D (green). Lines indicate the COM distances of NTD heterodimers (**a**, left one), heterodimeric R2-R2 lobes (**a**, left two), R2-R2 and LBD-LBD between two N1 (**c**, left) or two N2 (**c**, right). The open-closed and twisting-untwisting dihedral angles of NTD clamshell in N1 and N2 are labeled with the corresponding color to the subunits (**a**). Arrows indicate the conformational change from N1-N2A to N1-N2D di-receptors. **d, e,** Molecular dynamics simulation of N1-N2A (**e**) and N1-N2D (**d**) receptors in Gly-Glu bound state. Left panels show the ensemble plot for the trajectory of N2D- and N2A-NTDs. Middle and right panels show the histogram and the corresponding fitting (sum of Gaussians) of the open-closure, twist-untwist dihedral angle of individual NTD, Cα-Cα distance of two S238s or A263s in N2D and two K220s or L246s in N2A subunits, respectively. **f**, DTT induced current amplitude changes on receptors of WT N1 subunit incorporating WT N2D (0.24 ± 0.02), N2D^S238C^ (0.26 ± 0.03), N2D^A263C^ (0.24 ± 0.01), WT N2A (0.72 ± 0.21), N2A^K220C^ (1.13 ± 0.15), N2A^L246C^ (1.34 ± 0.19) subunits, n=6-19 oocytes. Top panels exhibit NTD tetrameric interfaces (rounded rectangle) formed by α5-α5 in N2D, and α5 together with α6 in N2A (PDB:6MMP, ref.^10^) di-receptors. **g**, DTT induced current amplitude changes on receptors of N1-N2D (0.24 ± 0.02), N1^E698C^-N2D (1.56 ± 0.17), N1-N2D^2A-NTD^ (0.71 ± 0.01), N1^E698C^-N2D^2A-NTD^ (0.70 ± 0.06), N1-N2A^2D-NTD^ (0.48 ± 0.17), N1^E698C^-N2A^2D-NTD^ (1.05 ± 0.20), N1-N2B^2D-NTD^ (0.60 ± 0.14), N1^E698C^-N2B^2D-NTD^ (1.10 ± 0.20), n=4-19 oocytes. Top panels are cartoon illustrations of WT and NTD plus NTD-LBD linkers switched chimeric receptors, the N2D, N2A and N2B subunits are colored in green, orange and violet, respectively. **h,i,** Single-channel recordings on proteoliposome incorporated with purified N1^E698C^-N2D receptors, holding at +20 mV. (**h**) shows the normalized all-point amplitude histogram and statistical analysis of Po. (**i**) shows the representative current traces. The distribution data were fitted by a sum of two Gaussians with two peaks corresponding to the close and open states. **j,** Top-down view of electron density of LBD layer in N1^E698C^-N2D receptor in crosslinked (marked as C-C) and non-crosslinked (marked as C-C) state. The density of E698C in N1 are highlighted in yellow mesh. All data shown are with mean ± SD; P values are determined by one-way ANOVA with Tukey’s multiple comparison test in (**f**) and by two-tailed unpaired Student’s *t*-test in (**g, h**) (***P<0.001; n.s means no significance).

By comparing these biologically-relevant structures, we noticed that the NTDs displayed similar configuration, especially with their tetrameric interfaces uniformly formed by α5 helices of N2D-NTDs (Fig.1h). By comparing Gly-Glu and Gly-CPP bound states, antagonist R-CPP binding to the N2D-LBD cleft opened the clamshell by 23.5° and subsequently pulled together D2-D2 lobes, leading to the shortened distance between two gate-linked α-helix E in LBD by 5.9 Å in N1 and 5 Å in N2D subunits and between two Cα of gate residues by 3.2 Å in N1 and 1.3 Å in N2D subunits, respectively (Fig.1i, j). Moreover, in the presence of exon 5, all four chains moved away from the central axis indicated by the expended of COM distance of two N1b- and two N2D-LBDs by 1.3 Å and 1.1 Å, respectively. And two α-helix E in N2D-LBDs that were pushed away by 1.9 Å in N1b-N2D receptor (Fig.1i, j). We suggest that this conformational movement triggered by exon 5 could account for more active gating properties in N1b-N2D receptors^5, 31^. These data implied that the molecular mechanisms of antagonist inhibition and splicing exon 5 modulation were conserved among the N1-N2D, N1-N2A^13^ and N1-N2B^12, 31^ di-receptors (Fig. 1k).

### Dominant negative role of N2D-NTD

As N1-N2A and N1-N2D di-receptors stand for two extremes of biophysical property in NMDA receptors^2^, we here compared their structures under the same condition of Gly-Glu bound state. Overall, the tetrameric NTD in N2D receptors underwent an anti-clockwise rotation and adopted a more compact configuration, as compared to that in N2A receptors (Fig.2a). Specifically, N2D-NTD intrinsically adopted a closer by 12° and more untwisted by 16.4° clamshell than N2A-NTD (Fig.2a). Molecular dynamics (MD) simulation further indicated that N2D-NTD was less mobile than the N2A-NTD, assessed by the angle measurement of twisting-untwisting and opening-closure (Fig.2d,e). This rigid conformation of N2D-NTD is in line with the previously-proposed functional studies^32, 33^. The configuration of N2D-NTD led to the separation of R2-R2 lobes within NTD heterodimer (Fig.2a) and “1-Knuckle” (solely formed by α5-α5 helices) conformation at the tetrameric interface in N1-N2D receptor (Fig.2d), compared with the “2-Knuckle” (composed of α5 and α6 helices) conformation in N1-N2A receptor^10^ (Fig.2e). Notably, our structure of N1-N2D receptor displayed similar NTD conformation as the zinc inhibited N1-N2A receptor trapped in the “1-Knuckle” state^10^ (Extended Data Fig.3a,b), explaining the reason why N2D-NTD has low affinity to zinc^34^.

Next, MD simulation further illustrated that the tetrameric interface was constantly formed by two α5 helices in N2D-NTDs (Fig. 2d). In two independent trajectories, the two α5 helices displayed closer contact indicated by Cα-Cα distance of two S238 residues, while two α6 showed consistent separation indicated by long Cα-Cα distance of two A263 residues (Fig.2d). To validate those structural observations, we introduced a cysteine substitution on N2D^S238^ in α5 helix (Fig.2f) and found a spontaneously-formed band corresponding to N2D-N2D homodimer in N1-N2D^S238C^ receptors (Extended Data Fig.3c,e). Two-electron voltage-clamp recordings (TEVC) on *xenopus* oocytes showed that N1-N2D^S238C^ receptors exhibited same relative Po (Extended Data Fig.3d) and current amplitude reduction induced by DTT treatment (Fig.2f), similar as WT N1-N2D receptors. As a control, N1-N2D^A263C^ receptors could not form disulfide crosslinking (Extended Data Fig.3e) and exhibited no impact on the channel activity (Fig.2f). In contrast, cysteine replacement on homologous sites of α5 and α6 on N2A subunit significantly boosted the channel activity after DTT reducing treatment, as compared to N1-N2A WT receptors (Fig.2e,f and Extended Data Fig.3f,g). Together with previous studies^32, 33^, we suggest that N2D-NTD is consistently stabilized in the untwisted and closed conformation, which has a dominant-negative impact on N1-N2D channel gating.

### Functional NTD-LBD cooperativity

Based on the structural comparison, NTD configuration was synchronously coupled to the downstream LBD layer (Fig.2b,c), with more constrained N1-LBDs and more expanded N2D-LBDs (Fig.2b,c) than that in N1-N2A receptors. To check whether crosslinking the LBD interface would affect the channel gating of N1-N2D receptors, we introduced individual cysteine substitution at the sites of V697, E698 and L699 on N1-LBD (Extended Data Fig.3h,i). The N1^C744A-C798A^-N2D receptors without endogenous redox sensor^35^ exhibited no change in current amplitude, while the N1^C744A-C798A-E698C^-N2D receptors showed the most markable inhibition by 2.6-fold after DTT treatment (Extended Data Fig.3i). Next, we estimated the relative channel activity by measuring the kinetics of MK-801 induced current inhibition^36, 37^ on N1-N2D receptors. N1^E698C^-N2D receptors displayed 10-fold faster inhibition kinetics than the wild-type (WT) receptors (Extended Data Fig.3j). Biochemical analysis further demonstrated that N1 subunits spontaneously formed homo-dimer in N1^E698C^-N2D receptors (Extended Data Fig.3l). In addition, Gly and Glu sensitivities of N1^E698C^-N2D receptors significantly declined by 170- and 4-fold, respectively (Extended Data Fig.3m). Together, we demonstrated that single cysteine replacement at E698 of N1 subunit dramatically switched the channel of N1-N2D receptors from a low- to high-activity state.

To explore whether NTD and LBD conformation is functionally coupled, we swapped NTD plus NTD-LBD linker between N2D and N2A or N2B subunits. As N1^E698C^-N2D receptors could be spontaneously locked in a super-active state (Fig.2g and Extended Data Fig.4j, k). Conversely, N1^E698C^-N2D^(2A-NTD)^ receptors lost current boosting effect in response to the DTT treatment and showed no significant difference as N1-N2D^(2A-NTD)^ receptors (Fig. 2g). Previous studies reported that N1^E698C^-N2A receptors decreased the channel activity by ∼2-fold, while the N1^E698C^-N2B receptors showed no significant effect to the reducing reagent, compared to the corresponding WT receptors^38, 39^. Strikingly, both N1^E698C^-N2A^(2D-NTD)^ and N1^E698C^-N2B^(2D-NTD)^ chimeric receptors showed significant current reinforcement in response to DTT reduction, in comparison with corresponding chimeric receptors with no cysteine substitution on N1^E698^ site (Fig.2g). These data confirm that the conformation of N2D-NTD allosterically controls the functional crosslinking of tetrameric LBDs.

### Functionally and structurally trapping N1^E698C^-N2D receptors

Next, we purified the protein of CTD-deleted N1^E698C^-N2D receptors and found a spontaneously crosslinked complex of ∼250 kDa corresponding to N1 homo-dimers (Extended Data Fig.4a, b). Then, we reconstructed the purified N1^E698C^-N2D protein into the proteoliposomes and performed single-channel recordings to evaluate the absolute Po. We found two independent channel activity profiles with one population displaying hyperactive (C-C state) Po of 0.226 ± 0.028 and the other (non C-C state) of 0.021 ± 0.002. The latter was close to the value previously found for WT N1-N2D receptors^5^ (Fig.2h,i). Based on these results, we concluded that our receptor in the C-C state was likely to be trapped in a hyperactive conformation.

We then determined the structure of N1^E698C^-N2D receptors in complex with Gly-Glu and found two classes of EM maps with different LBD conformations (Fig.2j and Extended Data Fig.4c, d). One structure at 6.4 Å resolution displayed two tethered N1-LBD protomers that were crosslinked at the E698C residue (named “C-C state”, Fig.2j). The other structure at 4.3 Å resolution with non-crosslinked N1-LBD exhibited a similar conformation as Gly-Glu bound WT receptors, with root mean squared deviation (r.m.s.d.) of 1.3 Å (“non C-C state”, Fig.2j). Structural comparison between Gly-Glu and C-C states revealed that disulfide crosslinking through N1^E698C^-N1^E698C^ glued together two N1-LBD and expanded two N2D-LBD protomers, as indicated by the center-of-mass (COM) distances (59.5 Å of Gly-Glu *vs* 56.3 Å of C-C for N1; 55.5 Å of Gly-Glu *vs* 65.1 Å of C-C for N2D, Extended Data Fig.4e). Strikingly, the two α-helix E in N2D-LBDs (directly linked to TM3) underwent a large outward movement, with their COM distance changed from 76.9 Å in Gly-Glu to 89.7 Å in C-C state. To date, among all the resolved NMDA receptor structures, our C-C state receptors exhibit the largest separation of D2-D2 lobes of two N2 subunits (Extended Data Fig.4f). Taken together, these functional and structural results are in line with each other, proving that a single disulfide crosslinking by GluN1^E698C^ at the tetrameric interface of LBD could convert low Po of N1-N2D receptors to high active state.

### Asymmetric structure of N1-N2C di-receptor

N1-N2C receptors display distinctive biophysical properties including rapid channel opening rate, no cluster bursting current and asymmetric transition between low and high conductance levels^4^. As no structural information is currently available, we purified the protein of CTD-truncated N1-N2C receptor (Extended Data Fig.5a, b) and elucidated its cryo-EM structures in the presence of Gly-Glu. After multiple rounds of 3D classification, the whole dataset yielded in three classes, with a major (87% population) and an intermediate classes (5.5% population) in asymmetric, and a minor class (7.5% population) in symmetric conformation. The asymmetric feature was mainly characterized by the geometry viewed from top-down NTDs (Extended Data Fig.5c). Therefore, we carried out the 3D refinement for the major and minor classes with C1 and C2 symmetry, respectively, and obtained two density maps of N1-N2C di-receptor at 3.6 Å and 4.3 Å resolution, respectively (Fig. 3b,c and Extended Data Fig.5c). Overall, the TMD signal in these structures was not well-resolved, which was presumably linked with the dynamic nature of TMD in N1-N2C di-receptor. To note, there was a subclass isolated from the major class displayed certain signal of TM2 and TM3 (Fig. 3a). To validate subunit identity, we performed mass spectrometry with the purified protein and confirmed 11 and 5 glycans on N1 and N2C subunits, respectively, in agreement with signals on the EM map for sugar moieties (Extended Data Fig.6). Therefore, we confirmed the N1-N2C di-receptor is globally assembled in a N1-2C-N1-2C pattern, with a dimer-of-dimer structure exhibiting domain swapping feature as other N1-N2 subtypes^6–11^ (Fig. 3a-c).

**Fig. 3.**
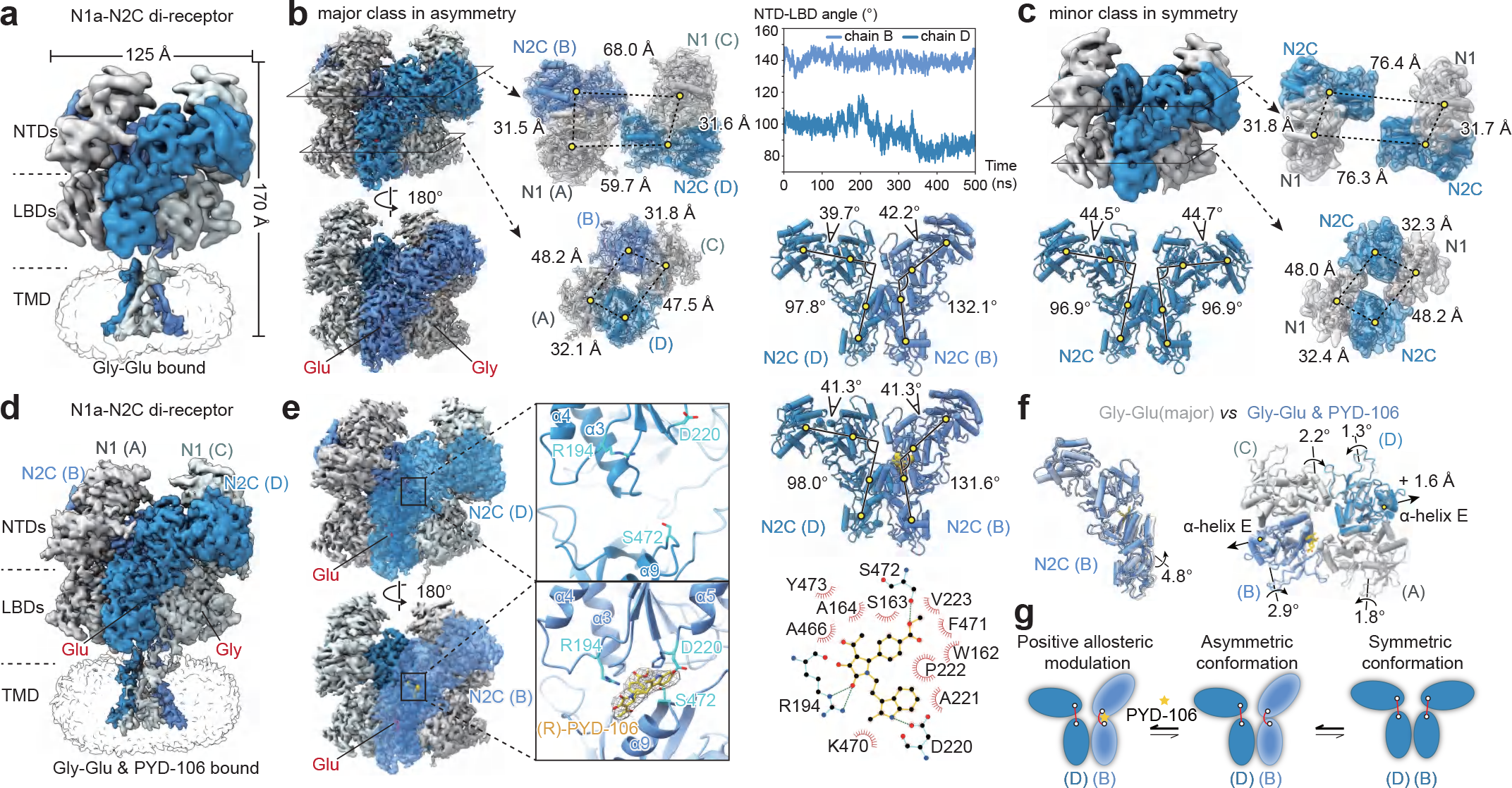
Cryo-EM structures and unique allosteric modulation of N1-N2C di-receptor. **a**, **d**, Intact cryo-EM structures of N1-N2C di-receptors in the Gly-Glu (**a**) and Gly-Glu and PYD-106 (**d**) bound state. Both maps are subclasses isolated from the respective major class, see Extended Data Figure 5,7. N1 and N2 subunits are marked as chain A/C and chain B/D, respectively. Agonists glycine and glutamate colored in red. **b**,**c** Structural analysis of N1-N2C di-receptors in the major asymmetric (**b**) and in the minor symmetric (**c**) conformation. Side-views of EM maps, top-down views of NTDs and LBDs tetramers are illustrated. Conformation comparison between two N2C subunits are indicated by vector angles connecting the COMs of R2 and R1, D1 and D2 lobes of N2C protomers. r.m.s.d trajectories for two N2C protomers (chain B and chain D) based on the asymmetric structure along the total simulation of 500 ns are shown in the middle panel. **e**, Structural analysis of N1-N2C di-receptors in Gly, Glu and PYD-106 bound state. The frontal N2C is set to transparent for clarity. Middle panel shows the zoom-in view of NTD-LBD interface in two N2C protomers. EM density of (R)-PYD-106 (in golden) at the NTD-LBD interface of chain B. Residues forming hydrogen bond interactions with PYD-106 are shown in cyan sticks. Right panel shows the conformation comparison between two N2C subunits and detailed interactions between PYD-106 and N2C analyzed by Ligplot^+^. **f**, Conformational change induced by positive allosteric modulation of PYD-106. Left panel shows NTD superimposed of N2C subunits (chain B in asymmetric class) of Gly-Glu (in white) and PYD-106 (in blue) bound structures, with LBD rotation angle indicated. Right panel shows the superimposition of LBD layers of Gly-Glu (major class) and PYD-106 bound state, with inward rolling degrees of four LBDs and distance changes in-between two α-helix E of N2C subunits indicated upon PYD-106 binding. **g**, Cartoon illustration for the dynamic transition of N1-N2C di-receptor among symmetrical(minor), asymmetrical(major) and PYD-106-bound conformations.

In the major class, the electron density of extracellular domains (ECDs) was in decent quality, and Gly and Glu were clearly visible within LBD clamshells of N1 and N2C, respectively (Fig.3b and Extended Data Fig.9a). Notably, both heterodimer subunits exhibited a largely asymmetrical configuration. Particularly on the NTD layer, inter-subunit COM distance between chain A and D (59.7 Å) was substantially smaller than that between chain B and C (68.0 Å) (Fig.3b). As the controls, the two N1- or N2C-NTD protomers and the two heterodimers exhibited conformational similarity (Extended Data Fig.9c). On the LBD layer, four LBDs exhibited a pseudo two-fold symmetry (Fig.3b and Extended Data Fig.9d). We attributed this transition from asymmetric NTD and symmetric LBD was due to different conformations of the NTD-LBD linker in two N2C subunits (Fig.3b and Fig.4d). In this major class, two N2C protomers of identical sequence exhibited distinct conformations, with the NTD-LBD angle of 132.1° in chain B and 97.8° in chain D. We carried out MD simulation on this asymmetric structure and found both N2C subunits exhibited stable asymmetric conformation in the respective to the NTD-LBD angles (Fig. 3b).

**Fig. 4.**
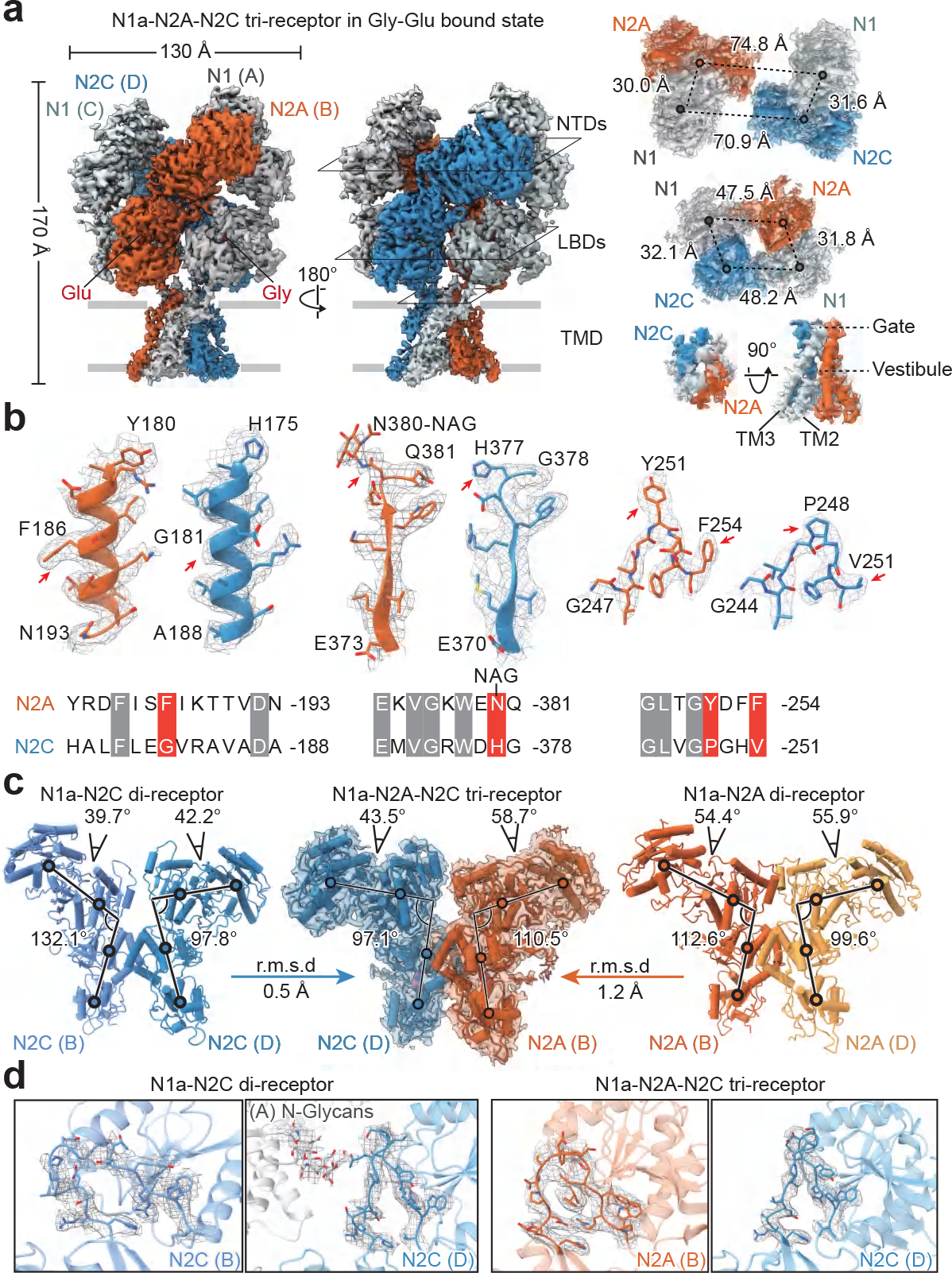
Structural analysis of N1-N2A-N2C tri-receptor. **a**, Cryo-EM structures of N1-N2A-N2C tri-receptor in Gly-Glu bound state. Top-down views of NTDs, LBDs and TMD are shown on the right. **b**, Local EM density illustration (top panel) and corresponding sequence alignment (bottom panel) of N2A and N2C subunits in the tri-receptor. Homologous residues between N2A and N2C subunits with the different EM densities are indicated by red arrows. The glycosylation density is marked at the site of N2A^N380^, in comparison to the non-glycosylated N2C^H377^ site. **c**, Conformation comparison of N2A and N2C in tri-receptor and respective di-receptors. NTD-LBD angles and NTD clamshell opening angles are all indicated. The r.m.s.d for ECD alignment between the N2A in tri- and di-receptor (chain B *vs* chain D) is 1.2 *vs* 2.2 Å, and N2C in tri- and di-receptor (chain B vs chain D) is 6.3 *vs* 0.5 Å, respectively. **d**, Zoom-in views of NTD-LBD linker density of N2 subunits in N1-N2C and N1-N2A-N2C receptors. The density for the NTD-LBD linkers in N1-N2A di-receptor (PDB: 6MMP) is not clear enough to show.

In the minor class, we found that both N2C subunits adopt the conformation of chain D in the major class, characterized by the NTD-LBD angle of 96.9° (Fig. 3c). On the NTD layer, the two NTD heterodimers largely sprayed from each other with inter-subunit COM distance in 76.4 Å (Fig. 3c). Moreover, the intermediate class exhibited a conformation situated between the major and minor classes. Aligning the maps among these three classes displayed that a rotation of one NTD heterodimer by 33.8°. MD showed that this symmetric conformation was highly mobile (with r.m.s.d ranging from ∼0.4 nm to more than 1.4 nm) and have the tendency to shit its conformation to the asymmetric state (Extended Data Fig.5d). Thus, we propose that the N1-N2C di-receptor can spontaneously transit between the symmetric and asymmetric conformations, with the asymmetric conformation more energetically favorable (Fig. 3g).

### N2C-specific positive allosteric modulation

A series of pyrrolidinones (PYDs) have been identified as positive allosteric modulators (PAMs) specifically for N1-N2C di-receptors with the potency in micromolar range^40^. Functional studies revealed that the potentiation effect of PYD-106 was induced by one enantiomer^40^ and was completely lost in the N1-N2A-N2C tri-receptors^41–43^. To date, the accurate binding site and allosteric mechanism of PYD-106 remain unknown. Hence, we tried to resolve the cryo-EM structure of N1-N2C receptor in the presence of Gly-Glu and PYD-106. Finally, a cryo-EM map at 3.0 Å resolution was obtained, with no symmetry applied (Fig.3d,e and Extended Data Fig.7). In our map, a clear electron density was only found at the compact NTD-LBD interface of N2C subunit in chain B, but not in chain D (Fig.3e), which was not predicted by previous functional and modeling data^41, 43^. The enantiomer of (*R*)-PYD-106 could perfectly fit into the geometry of electron density, while (*S*)-PYD-106 was stoichiometrically unfavorable (Extended Data Fig.10a). Ligplot^+^ analysis showed that three pairs of hydrogen bond interactions, between Arg194 and carbonyl on pyrrolidinone, Asp220 and nitrogen of the indole ring, and Ser472 and methyl ester. These interactions stabilized the three aromatic nuclei of PYD-106 (Fig.3e), in line with electrophysiological findings that mutation at any of above-mentioned residues resulted in the complete loss of PYD-106 potentiation^41, 43^. Moreover, residues at the bottom of R2 lobe (especially Pro222) and top of D1 lobe (Ala466 to Tyr473) formed hydrophobic interactions with PYD-106 (Fig.3e).

The binding of PYD-106 resulted in a 4.8° rotation of LBD relative to NTD in N2C of chain B and generated asymmetrical inward rolling of all four LBD clamshells (Fig.3f). Consequently, the allosteric binding of PYD-106 led to conformational changes in N2C-LBD, reflected by the 1.6 Å outward movement of α-helix E, which directly linked to channel gate (Fig.3f and Supplementary Video 1). To examine the basis of PYD-106 selectivity for N1-N2C di-receptors, we performed sequence alignment and found that Arg194, Asp220 and Ser472 in N2C showed low conservation with homologous residues in N2A/2B/2D subunits (Extended Data Fig.10b). Moreover, structural comparison showed that only in the chain B of N1-N2C receptors, the NTD-LBD interface formed adequate R2-D1 contact area (dSASA of 908.4 Å^2^), yielding the lowest interface binding energy (dG_separated of -7.7 REU), as compared with that of other N2 subunits (Extended Data Fig.10c). Notably, the two N2C protomers also adopted asymmetric conformation with NTD-LBD angles of 131.6° in chain B and 98.0° in chain D (Fig.3e), consistent with the asymmetric conformation in the major class of Gly-Glu bound state (Fig.3b). In summary, our data indicates that PYD-106 preferably locked the N1-2C di-receptor in the asymmetric conformation (Fig. 3g).

### Architecture of triheteromeric N1-N2A-N2C receptor

Previous studies have proposed that N1-N2A-N2C tri-receptors are the dominant population in cerebellar granule cells, and these tri-receptors retain biophysical properties of both N1-N2A and N1-N2C di-receptors^19, 42, 44^. In order to explore its architecture, we fused His- and Strep-tags on N2A and N2C constructs respectively, and isolated tri-receptors by two-step affinity chromatography. Western blotting analysis verified the presence of both N2A and N2C subunits in purified tri-receptor proteins (Extended Data Fig.8a,b). Eventually, we obtained a 3.5 Å resolution EM density map of N1-N2A-N2C receptor in complex with Gly-Glu (Fig.4a and Extended Data Fig.8c, 9a,b). Mass spectrometry revealed 6 and 5 glycans on N2A and N2C subunits, respectively (Extended Data Fig.6). Glycans presence at N2A^N380^ position and absence at the homologous site of N2C^H377^ could clearly distinguish N2A from N2C subunits. Furthermore, our high-resolution map also provided clear signals of side chains to authenticate N2A and N2C subunits, based on non-conserved NTD sequences (Fig.4b). Taken together, we confirmed that this tri-receptor assembled in a N1-2A-N1-2C arrangement and adopted an asymmetric architecture on NTDs, LBDs and TMD layers (Fig.4a).

Based on the knowledge that both N1-N2A^10, 45^ and N1-N2C di-receptors exhibited asymmetric features, it is of interest to note that N2A and N2C subunits in the tri-receptor share similar conformation with that of chain B in N1-N2A (PDB: 6MMP, ref.^10^) and of chain D in N1-N2C di-receptors, respectively, as characterized by NTD-LBD angles and ECD superimposition (Fig.4c). We also stretched the linker conformation from our high-resolution structures of N1-2C di- and N1-2A-2C tri-receptors, and found both N2C linkers in chain D adopted similar vertical configuration, while N2C in chain B of di-receptors adopted a flattened configuration (Fig.4d). Furthermore, this asymmetric arrangement was adopted in the LBD layer as well. Superimposing the N1-LBDs within the intra- or inter-dimer, we found that N2A-LBD displayed a 4.3° rotation or 4.8° inward rolling, relative to the conformation of N2C-LBD (Extended Data Fig.9d). These findings demonstrate that one protomer conformation of N2A and N2C in the corresponding di-receptors was precisely integrated into the N1-N2A-N2C tri-receptors (Supplementary Video 2).

Therefore, the PYD-106 insensitivity of N1-N2A-N2C tri-receptor could be attributed to the fact that N2C in the tri-receptor adopted the conformation of chain D in N1-N2C di-receptor, with R2-D1 lobes spatially disconnected (Extended Data Fig.10c). The structural and functional results indicate that the PYD-106 binding highly depends on the conformational specificity of N2C subunit adopted in di- or tri-receptors. Altogether, these findings provide the structural insight into the distinct biophysical and pharmacological property of the tri-receptors^42^.

## Discussion

In this work, we uncovered molecular architecture and assembling feature in N2C- or N2D-containing NMDA receptors, and provided structural insights into the subtype-specific gating mechanism and pharmacological property of these subtypes. In order to elucidate the molecular determinants for distinct biophysical and pharmacological profiles^2, 15^, we performed a comprehensive structural analysis of major NMDA subtypes, including N1-N2A-N2C tri-receptor and four types of N1-N2 di-receptors in the same Gly-Glu bound state (Fig.5a). On the NTD layer, from N2A to N2D di-receptors, individual N2-NTD adopted progressively open to close conformation, and R2-R2 distance within the NTD heterodimer also increased gradually (Fig.5b). At the tetrameric interface, the separation of two R2 lobes in N2-NTD increased gradually from N2A to N2D di-receptors (Fig.5b). These results strongly support the notion that differential function of NMDA receptors is controlled by the conformation of N2-NTD^32, 33^.

**Fig. 5.**
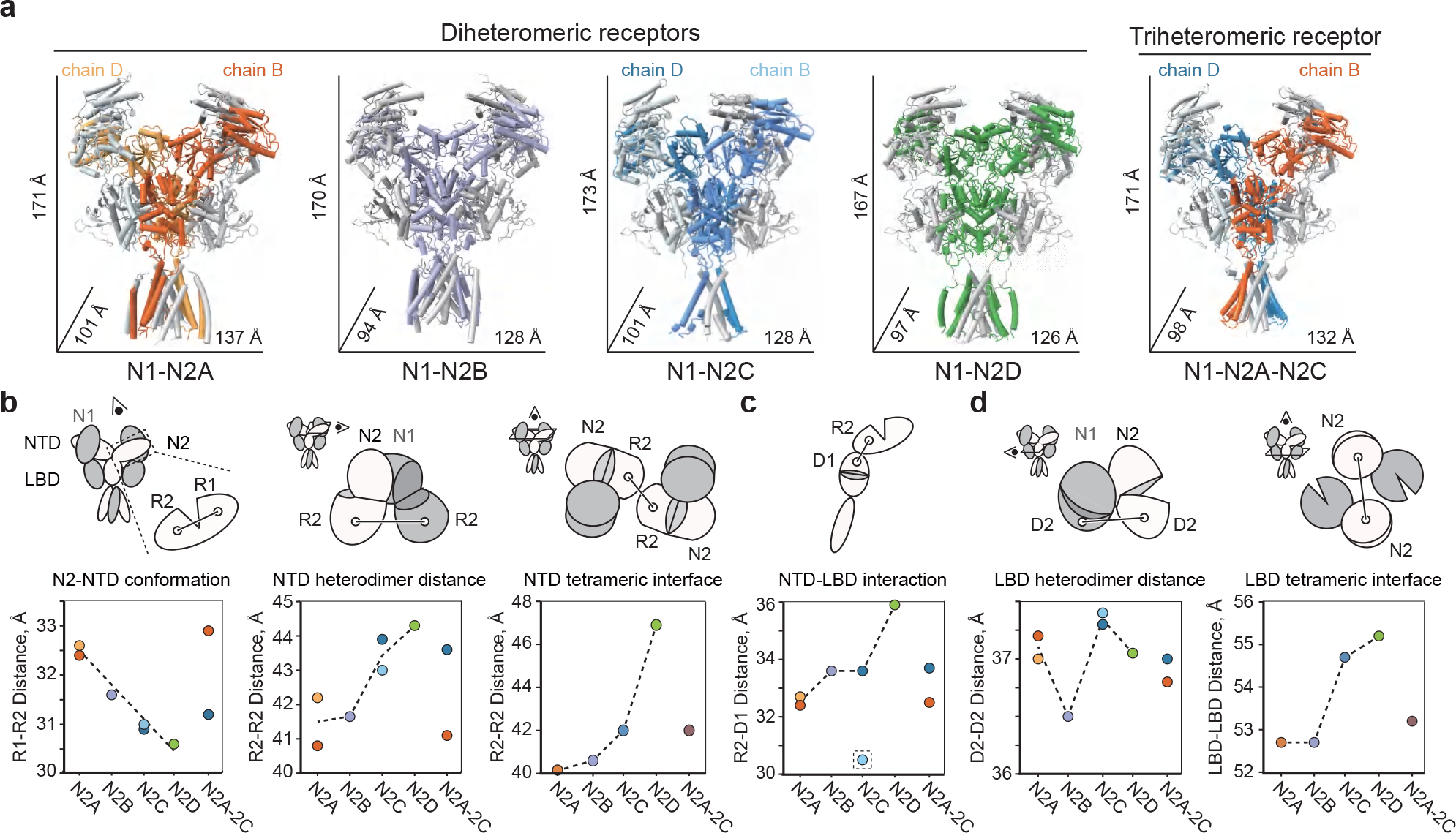
Comprehensive structural analysis of diverse NMDA receptors. **a**, A gallery of cryo-EM structures of N1-N2A (PDB: 6MMP, ref.^10^), N1-N2B (PDB: 7EU8, ref.^14^), N1-N2C and N1-N2D di- and N1-N2A-N2C tri-receptors at Gly-Glu bound state. **b-d**, Scatter diagram of COM distance in various NMDA receptor subtypes. Schematic representation of the tetrameric NMDA receptor with N1 colored in grey and N2 in white, respectively. Six parameters measuring the distance between COMs of lobes and domains are indicated. For the asymmetric N1-N2A, N1-N2C di- and N1-N2A-N2C tri-receptors, two N2 subunits of chain B and chain D with different geometry were individually measured. NTD conformations of monomer, heterodimer and tetramer are indicated by COM distances of R1-R2 in N2, R2-R2 in heterodimer and R2-R2 between two N2-NTDs, respectively (**b**). NTD-LBD conformation is characterized by R2-D1 distance in the N2 subunit, with dashed box indicating the chain B of N2C subunit in N1-N2C di-receptors (**c**). LBD conformations of dimer and tetramer are indicated by COM distances of D2-D2 in heterodimer and two N2-LBDs, respectively (**d**). Dashed lines connecting parameters from N2A to N2D di-receptors show the tendency of conformational transition among these four major subtypes.

To evaluate the subunit cooperativity between NTD and LBD layers, we noticed that N2A subunits exhibited strong interactions while N2D subunits displayed loose contacts, as shown by the distance between NTD-R2 and LBD-D1 lobes (Fig.5c). In the structure of N1-N2A-N2B tri-receptors, N2A was also shown to adopt a more extensive NTD-LBD interaction than N2B subunit^46^. As an exception, the chain B of N2C in di-receptors displayed the most extensive interaction between R2 and D1 lobes (Fig.5c, dashed box). This was further evidenced by the asymmetric assembly and unique PYD-106 binding feature in N2C di-receptors (Fig.3).

On the LBD layer, neither individual clamshell of N2 (with D1-D2 distance ∼ 26.5 Å) nor N1-N2 LBD heterodimer (D2-D2 distance in Fig.5d) showed consistent conformational tendency among four subtypes of di-receptors. With respect to the LBD tetramer, two N2-LBDs increasing moved apart from N2A to N2D di-receptors. This is likely to reflect allosteric coupling with the tendency of increasing separation between the two R2 lobes on NTD layer (Fig.5d). Finally, in all above measured parameters, the individual N2A and N2C subunits in tri-receptors retained structural features similar as the corresponding N2A (chain B) and N2C (chain D) chains in both di-receptors (Fig.5b-d). During this paper revision, we noticed that a very recent work from Furukawa’s lab also reported the structural insights into assembly and function of N1-N2C, N1-N2A-N2C and N1-N2D NMDA receptors^47^. Altogether, these findings revealed the molecular structure and functional diversity of NMDA receptors and pave the way for the structure-based drug design with subtype specificity.

## Data availability

Cryo-EM density maps of Gly-Glu bound N1a-N2D, Gly-CPP bound N1a-N2D, crosslinked N1a^E698C^-N2D, non-crosslinked N1a^E698C^-N2D, Gly-Glu bound N1b-N2D, Gly-Glu bound N1a-N2C in asymmetric and symmetric conformation, Gly-Glu & PYD-106 bound N1a-GluN2C and Gly-Glu bound N1a-N2A-N2C receptors have been deposited in the Electron Microscopy Database under the accession codes EMD-33792, EMD-33788, EMD-33795, EMD-33798, EMD-33793, EMD-33789, EMD-34674, EMD-33790, EMD-33791, respectively. The coordinates for the structures have been deposited in the PDB under accession codes 7YFL, 7YFF, 7YFO, 7YFR, 7YFM, 7YFG, 8HDK, 7YFH and 7YFI, respectively. Additional data that support the findings of this study are available from the corresponding author upon request.

## Acknowledgements

We thank Dr. Yu Kong and Lijun Pan at Electron Microscopy Facilities of Center for Excellence in Brain Science and Technology, Chinese Academy of Sciences for assistance with sample screen. We thank Dr. Boling Zhu and Dr. Xujing Li at Center for Biological Imaging (CBI), Institute of Biophysics, Chinese Academy of Sciences, Dr. Qingxia Wang and Yue Zhou at Electron Microscopy Facility, Shanghai Institute of Materia Medica (SIMM), Chinese Academy of Sciences, Dr. Qianqian Sun and Yaping Wang at Bio-Electron Microscopy Facility, ShanghaiTech University for their help in Cryo-EM data collection. We thank Dr.Yanyan Jia at Fudan University for the assistant on single-channel recordings. We are grateful to Dr. Muming Poo for proofreading. Financial supports are gratefully acknowledged for the National Science and Technology Innovation 2030 Major Program (2022ZD0212700), the Lingang Laboratory (LG 202106-02), the China State Key Research Grant (2017YFA0505700), the Strategic Priority Research Program of Chinese Academy of Science (XDBS01020000) and the Shanghai Municipal Science and Technology Major Project (2018SHZDZX05).

## Author contributions

J.Z. and M.Z. purified and froze the protein, collected and analyzed the cryo-EM data, built atomic model and conducted electrophysiology of N2D and N2C receptors, respectively; H.W., Q.W. and Y.W. carried out *in silico* calculations; Z.L. and F.W. performed mass spectrometry; N.S. and Z.K. performed biochemistry assays; F.Y. carried out single-channel recordings; Y.L. and F.G. participated in data collection of N2C receptors; J.Z., M.Z. and S.Z. wrote the manuscript. S.Z. conceived the project and supervised the research.

## Methods

### Plasmid construction

For structural elucidation of N1-N2D receptors, the cDNAs of wild-type (WT) *Homo sapiens GRIN1-1a* (M1-Q847, NM_007327.4) *GRIN1-1b (*M1-Q868, NM_001185090.2) and *GRIN2D* (M1-P879, NM_000836.4) were cloned into pEG-BacMam vector^1^. Strep-tag II was fused to the C terminus of N2D. For biochemistry and electrophysiology, full-length WT *Homo sapiens GRIN1-1a*, *GRIN2D, GRIN2A* (NM_001134407.3)*, GRIN2B* (NM_007327.4) were cloned into pCI-neo vector^2^, a gift from Prof.Hongjie Yuan (Emory University, USA). Chimeric constructs were generated by the Gibson assembling kit (NEBuilder® M5520AA). Mutant constructs were obtained by performing site-directed mutagenesis on WT constructs using Takara KOD-FX DNA polymerase. For structural studies of N2C-containing receptors, wild-type *Rattus norvegicus GRIN1-1a* (M1-Q847, NP_058706), *GRIN2A* (M1-F841, NP_036705) and *GRIN2C* (M1-V839, NP_036707) followed by a 3C protease cleavage site (LEVLFQGP) were also cloned into pEG-BacMam vectors. For N1-N2C receptor, a 6×His tag, an mRuby encoding sequence followed with a Strep-tag II were fused to the C terminus for N1 and N2C, respectively. For N1-N2A-N2C receptor, a GFP sequence followed with an affinity tag of 6×His or Strep-tag II were placed at the C terminus of N2A or N2C. N2A and N2C subunits were subcloned into one vector linked by T2A (EGRGSLLTCGDVEENPGP) self-cleaving peptide.

### Protein expression and purification

Recombinant baculovirus was produced using sf9 insect cells following the instructions of Bac-to-Bac TOPO Expression System (Invitrogen A11339). Suspended HEK293S GnTI^-^ cells at the concentration of 3.5-4.0×10^6^/mL in 37°C were infected with P2 virus. 8-12 hours post-infection, 10 μM MK-801 and 10 mM sodium butyrate were co-added into the culture medium for boosting protein expression. Cells were transfer to 30°C for additional 48-60 hours, and then collected by centrifugation at 7,000 g for 20 min.

Cell pellet was resuspended and sonicated with TBS (150 mM NaCl, 20 mM Tris, pH 8.0), solubilized in TBS buffer supplemented with 1% lauryl maltose neopentyl glycol (L-MNG), 2 mM cholesterol hemisuccinate (CHS), a protease inhibitor cocktail of 0.8 μM aprotinin, 2 mM pepstatin A, 2 μg/mL leupeptin and 1 mM phenylmethyl sulfonyl fluoride (PMSF), 1 mM Gly, 1 mM Glu and 100 μM ethylenediamine tetraacetic acid (EDTA) for 1.5 h at 4°C. After ultracentrifugation at 40,000 rpm, the supernatant was incubated with streptactin resin. The resin was rinsed with wash buffer (TBS supplemented with 0.1% L-MNG, 2 mM CHS, 2 mM Gly, 2 mM Glu and 100 μM EDTA), and eluted with wash buffer supplemented with 5 mM D-desthiobiotin. The eluted protein was concentrated and further injected into a Superose 6 Increase column (GE Healthcare) for size-exclusion chromatography (SEC) in TBS buffer supplemented with 0.1% digitonin, 5 μM CHS, 0.1 mM CHAPSO, 1 mM Gly, 1 mM Glu and 100 μM EDTA. The peak fraction was pooled and concentrated to approximately 4 mg/mL for cryo-EM grid preparation.

For N1-N2A-N2C tri-receptor, 5 mM imidazole was added to the elution from streptactin affinity chromatography, and the sample was bound to IMAC nickel resin for His-tag purification. Nickel resin was rinsed with wash buffer supplemented with 15 mM imidazole, and the tri-receptor protein was eluted using wash buffer supplemented with 250 mM imidazole. 400 μM (*R*/*S*)-PYD-106 (AOBIOUS, AOB6695) was added into N1-N2C protein for the PYD-106 bound structure, and 1 mM Gly and 600 μM (*R*/*S*)-CPP (SIGMA, C104-25MG) were added into N1a-N2D protein for Gly-CPP bound structure. All purification procedures were conducted at 4°C.

### Cryo-EM sample preparation, data acquisition and processing

3 μL protein sample was applied to the glow-discharged 300 mesh Quantifoil R1.2/1.3 (Au) grid (ELECTRON MICROSCOPY CHINA, BQR1.2/1.3-3A) using FEI Vitrobot (Thermo Fisher), with the chamber environment controlled at 8 °C and in 100% humidity. Grids were blotted for 3 s and immediately plunged into liquid ethane for vitrification.

Cryo-EM data were collected on 300 kV Titan Krios G3 electron microscope (FEI) equipped with K3 Summit direct electron detector (Gatan) and GIF quantum energy filter. The pixel size in the super-resolution counting mode was set to 0.5355 Å for all datasets of N2C-containing receptors and to 0.415, 0.5335, 0.5355 Å for different datasets of N2D-containing receptors. Each movie stack was dose-fractionated over 40-50 frames with a total dose of 60 electrons. The defocus values of movie stacks varied between –1.2 μm and –2.5 μm. Automatic data collection was conducted using Serial EM.

Beam-induced motion and drift correction were performed using MotionCor2,ref^3^. CTF parameters for each micrograph were determined by Gctf^4^. Approximately 2,000 particles were manually picked and subjected to an initial reference-free 2D classification. Auto-picked particles were extracted and then subjected to several rounds of reference-free 2D classification and 3D classification. For N2C-containing receptors, no symmetry was applied during the data processing. For N1-N2D receptors, C2 symmetry was applied for the 3D refinement procedure. The ‘gold-standard’ FSC resolution were calculated with a soft shape mask applied to independent unfiltered half maps, with 0.143 criteria^5^. Data processing were mainly conducted by Relion 3.1.1^6^, except for 3D classification and 3D refinement in the datasets of Gly-Glu and Gly-CPP bound N1-N2D receptors were conducted by CryoSPARC^7^. More detailed information of data collection and processing are available in Extended Data Fig.1, 4, 5, 7 and Extended Data Table 1.

### Model building and refinement

Initial templates of N1a-N2D, N1b-N2D, N1-N2C di-receptors and N1-N2A-N2C tri-receptor were generated with SWISS-MODEL^8^, based on the homology structures of N1b-N2B receptor (PDB:6WI1, ref^9^) and N1a-N2A receptor (PDB:6MMT, ref^10^), respectively. Rigid-body docking with ChimeraX^11^ was used to fit the structural coordinates into the density maps. Flexible fitting was done with molecular dynamics flexible fitting (MDFF)^12, 13^ simulation with CHARMM36m force field^14^. The simulation temperature was maintained at 300 K using the Langevin algorithm^15^ and the Generalized-Born implicit solvent model^16, 17^ was used to describe the solvation effects. The models were then subjected to iterative manual adjustment in Coot 0.9.6.1^18^ and real space refinement in Phenix 1.20.1^19^. Local resolution of density maps is estimated by using ResMap-1.1.4^20^.

### Two-electrode voltage-clamp recording

*Xenopus* oocytes were prepared, injected and voltage-clamped as previously reported^21^. TEVC recording was performed in the extracellular solution containing (in mM) 100 NaCl, 2.5 KCl, 0.3 BaCl_2_, 5 HEPES, 0.01 DTPA, pH adjusted to 7.3 with NaOH. The maximum current of NMDA receptors was induced by co-application of 100 µM Gly and 100 µM Glu, at a holding potential of –60 mV. Current responses were recorded by pClamp 10 (Molecular Devices) and analyzed by Clampfit 10.6 software. In the DTT reduction experiment, oocytes were incubated for 15 min in solution containing (in mM) 5 DTT, 88 NaCl, 10 HEPES, 1 KCl, 2.4 NaHCO_3_, 0.33 Ca(NO_3_)_2_, 0.41 CaCl_2_, 0.82 MgSO_4_, pH adjusted to 7.6 with NaOH.

In order to evaluate the relative Po, 200 nM MK-801(Abcam, ab120027) was applied upon the maximum activation of the NMDA receptors. MK-801 inhibition constants (τ_on_) were calculated by fitting current response trace in inhibition phase (between 10%–90% of the maximal inhibition) with a single-exponential fit. Each value of mutant receptors was normalized to the mean τ_on_ of WT N1-N2D receptors.

### Reconstitution of proteoliposomes

Membrane protein was reconstituted into artificial liposomes as previously described^22^. Asolectin (Sigma-Aldrich, 11145-50G) was dissolved in chloroform at a stock concentration of 25 mg/mL and dried under liquid nitrogen stream to form a thin layer along the inner wall of glass tube. Subsequently, 1 mL 0.4 M sucrose was added and the tube was incubated at 50 °C in water-bath for 3 hours until the lipid cloud was formed. Purified CTD-truncated N1^E698C^-N2D receptor protein was then added at the weight ratio of protein: lipid (1:1000), and further incubated for 3 hours at 4 °C with a shaking speed at 100 rpm. Freshly reconstituted proteoliposomes were mixed with bath buffer containing (in mM) 140 NaCl, 2.8 KCl, 0.5 CaCl_2_, 10 HEPES and 0.01 BAPTA, pH adjusted to 7.3 with NaOH, and used for single-channel recording experiment.

### Single-channel recording

Single-channel recording was performed using pipette buffer containing (in mM) 140 CsCl, 5 EGTA, 0.1 Gly, 1 Glu and 10 HEPES, pH adjusted to 7.2 with CsOH, and above-mentioned bath buffer. Borosilicate glass pipettes (BF150-86-10, Sutter Instrument) were fabricated using P-97 puller (Sutter Instrument) with resistance in the range of 3-6 MΩ. The patch resistance increased to a gigaohm seal (>1GΩ) after the pipette formed a tight seal with the liposome membrane, single-channel currents were recorded with cell-attached patches at a holding potential of +20 mV and the data were collected with EPC-10 amplifier and Pulse software (HEKA Electronic, Lambrecht, Germany) with a 0.5-kHz low-pass filter and 50-Hz notch filter. 10 s duration from independent recordings was selected for data analysis. Open-shut state histogram was fitted with two exponential components.

### Western blotting

WT and mutant N1-N2D receptors were expressed in HEK293S cells by Polyethylenimine transfection (at the ratio of 1:3 of DNA and Polyethylenimine). Cells were collected and solubilized in a lysis buffer (20 mM Tris pH 8.0, 150 mM NaCl, 1% L-MNG, 0.8 μM aprotinin, 2 mM pepstatin A, 2 μg/mL leupeptin and 1 mM PMSF) for 1 h. Samples of N1-N2D or purified N1-N2A-N2C protein were subjected to loading buffer in the presence or absence of 0.5 M DTT and separated on SDS-polyacrylamide gel electrophoresis (4-6%). Protein were transferred to polyvinylidene fluoride membranes and immunoblotted with monoclonal antibodies at ratio of 1:1000 (anti-N1, Millipore, MAB1586; anti-N2D, Millipore, MAB5578; anti-strep, Abcam, ab252885; anti-6×his, Abcam, ab15149) and subsequently with secondary antibodies of HRP-conjugated (mouse IgG, Cell Signaling Technology, 7076s; rabbit IgG, Cell Signaling Technology, 7074P2). Protein bands were visualized by ECL substrate (Tanon, #180-501).

### MD simulation

Atomistic system of N1-N2A (7EU7^23^), N1-N2D, N1-N2C (both symmetric and asymmetric structures) embedded in 1-palmitoyl-2-oleoyl-sn-glycero-3-phosphocholine (POPC) bilayer was set up with CHARMM-GUI^24, 25^. One copy of protein and 771 POPC molecules were placed in a cubic simulation box. The system was solvated in 0.15 M NaCl solution, which makes up a total of 597467 atoms. Simulations were performed using GROMACS^26^ version 2021.3 with the CHARMM36 force field^27, 28^ and TIP3P water model^29^. The system was equilibrated in six steps with gradually decreasing restraining force constants on the protein. Gradually reduced constraints were applied during the equilibration. Two repeats of 200 ns unrestrained atomistic simulations were then performed. The Particle Mesh Ewald (PME) method^30^ was used to model the long-range electrostatics (< 1 nm). Temperature coupling was done with V-rescale thermostat^31^ at 310 K. The Parrinello-Rahman barostat^32^ with a reference pressure of 1 bar and a compressibility of 4.5e-5 bar was applied for pressure control. Covalent bonds are constrained to their equilibrium length by the LINCS algorithm^33^. The integration steps of all simulations were set to 2 fs.

### Mass spectrometry analysis

20 μg protein of N1-N2C or N1-N2A-N2C receptors was buffer-exchanged into denaturing buffer with 8 M urea, 50 mM NH_4_HCO_3_, pH 8.0. Samples were reduced and alkylated by tris(2-carboxyethyl) phosphine (TCEP) and Iodoacetamide (IAA), and exchanged into digestion buffer (10 mM NH_4_HCO_3_, pH 8.0) and digested by Lys-C at 37°C for 1h. Then, trypsin, Glu-C or Chymotrypsin was added respectively and further digested overnight. Supernatant was collected and acidified by 10% formic acid (FA) to pH 2.0 and were subjected to lyophilization and stored at -20°C until use.

The peptides were resuspended in 0.1% FA and 400 ng were injected for LC-MS/MS analysis. The peptides were first loaded onto a C18 trap column (3 μm, 120 Å) with Buffer A (0.1% FA in H_2_O) and then separated with a C18 analytical column (2.4 μm, 120 Å) by gradient elution of 5% - 35% buffer B (0.1% FA in acetonitrile) in 65 min. MS data were collected by Thermo Fusion Lumos mass spectrometer (Thermo Fisher Scientific) in a data-dependent acquisition (DDA) manner. The full MS spectra were obtained by Orbitrap analyzer with a resolution of 120,000 and a scan range of 350-1,800 m/z. Precursor ions with a charge state of 2+ to 8+ and a minimum intensity of 5E4 were isolated and subjected to high-energy collisional dissociation (HCD) with a normalized energy of 28% and electron-transfer/high-energy collision dissociation (EThcD). The calibrated charge-dependent ETD parameters were used and the SA collision energy were set at 15%. MS2 data including HCD and EThcD were obtained by Orbitrap analyzer with a resolution of 30,000.

MS data were all processed by pGlyco 3.0^34^. Protein sequences of sample were used as database for peptide searching. The max miss cleavage was set as 3. The mass tolerances of MS1 and MS2 were set at 10 and 20 ppm, respectively. The result were further analyzed and visualized using R with ’data.table’, ’tidyverse’ and ’ggsci’ packages.

### Structural analysis

For the structural measurements, NTD and LBD were separated into R1 and R2, D1 and D2 lobes, respectively. Center of mass (COM) of each domain or lobe was calculated in Pymol using “center_of_mass” script. NTD-LBD vector angers were calculated in Pymol using “vector_angle” script, with one vector connecting the COMs of NTD-R2 and NTD-R1 lobes, the other vector connecting the COMs of LBD-D1 and LBD-D2 lobes. Rotation between the domains was calculated using “draw_rotation_axis” script in Pymol. The dimension of receptors was obtained by running “draw_protein_dimensions” script in Pymol. Dihedral angles were measured by connecting the Cα atoms of indicated residues.

### Data analysis and statistic

For single channel recording, the Pulse files were converted into PCLAMP format using the ABF File Utility, Ver. 2.1.75 (Synaptosoft, Decatur, GA). Current traces of both single channel and TEVC recording were analyzed and fitted by Clampfit 10.7 (Axon Instruments, Foster City, CA). The GraphPad Prism 8.0 software (Graphpad Software, Inc.) was used for statistical analysis and graph generation.

