## Supplemental information for "Distinct structure and gating mechanism in diverse NMDA receptors with GluN2C and GluN2D subunits"

6.There authors contributed equally.

This file contains 10 Extended Data Figures, 2 Supplementary Videos and 2 Extended Data Tables.

**Extended Data Fig. 1 | Protein purification and cryo-EM analysis of Gly-Glu and Gly-CPP bound N1a-N2D receptors**

**a,** Schematic illustration of CTD-truncated receptor composed of human N1a (in grey) and strep tag fused N2D (in green) subunits.

**b-d**, Coomassie blue gel staining (**b**) and fluorescence SEC (FSEC) analysis of purified Gly-Glu (**c**) and Gly-CPP (**d**) bound N1a-N2D receptors protein.

**e, f**, Cryo-EM data-processing workflow of the Gly-Glu (**e**) and Gly-CPP (**f**) bound N1a-N2D receptors. Representative micrographs and 2D average classes are shown. Note that the special 2D averaged class with splayed extracellular domains (highlighted by a red box) was found in both datasets. For both datasets, motion correction, CTF estimation, particle picking and 2D classification were carried out by Relion 3.1.1[1]. Then, 3D classification and 3D refinement were subsequently performed by CryoSPARC[2].

**g, h**, Local resolution representation of density map and Fourier shell correlation (FSC) of Gly-Glu (**g**) and Gly-CPP (**h**) bound N1a-N2D receptor structures. Maps are colored based on the local resolution estimation by ResMap-1.1.4[3]. Masked (blue) and unmasked (green) FSCs of corresponding maps are both shown, where gold standard FSC = 0.143 was applied for the indication of final resolution (dashed line).

**Extended Data Fig. 2 | Protein purification and cryo-EM analysis of N1b-N2D receptors**

**a,** Schematic illustration of CTD-truncated receptor composed of human N1b (in grey) and strep tag fused N2D (in green) subunits.

**b**,**c,** Coomassie blue gel staining and FSEC analysis of purified proteins of N1b-N2D receptors.

**d,** Cryo-EM data-processing workflow of N1b-N2D receptor datasets processed by Relion 3.1.1[1]. Representative micrographs, 2D average classes, 3D classes and final density maps are shown.

**e,** Local resolution representation of density map and Fourier shell correlation (FSC) of Gly-Glu bound N1b-N2D receptor. Maps are colored based on the local resolution estimation by ResMap-1.1.4[3]. Masked (blue) and unmasked (green) FSCs of corresponding maps are both shown, with gold standard FSC of 0.143 criteria indicated.

**Extended Data Fig. 3 | Structural and functional analysis of N1a-N2D receptors**

**a**,**b,** Conformation comparison of Gly-Glu bound N1a-N2D and zinc-bound N1a-N2A (PDB:6MMK, ref[4]) di-receptors, as shown by NTD tetramer and heterodimer (**a,** left two), R1 lobes superimposed N1-NTDs or N2-NTDs (**a,** right two), and two N1 and N2 protomers (**b**). The open-closed and twisted-untwisted dihedral angles of N1-NTD and N2-NTD clamshells are labelled (**a**). Lines indicate the COM distances of domains or lobes.

**c**, Cartoon representation of NTD tetrameric interface in Gly-Glu bound N1a-N2D receptor structure. Residues of V231, Q235, S238 and A263 in N2D subunits are shown in sticks, with Cα-Cα distances indicated.

**d**, MK-801 inhibition kinetics on WT N1a-N2D and N1a-N2D$^{S238C}$ receptors. Left panel shows the recording traces of WT or mutant receptors in response to 200 nM MK-801 upon Gly and Glu (100 μM each) co-application. Recording traces were scaled to the maximum current amplitude induced by saturating agonists. Right panel shows the statistics of relative MK-801 inhibition off-rate kinetics ($\tau_{on}$, monoexponential fits) constants of N1a-N2D(1.00 ± 0.08) and N1a-N2DS238C(1.01 ± 0.08), n=10-12 oocytes. All values were normalized to the mean value of WT N1a-N2D receptors.

**e**, Western blotting analysis on WT and cysteine-substituted N1-N2D (**e**) and N1-N2A (**f**) receptors. Bands of N1 momomer, N2 momomer and N2-N2 homodimer are indicated.

**f**, Cartoon representation of NTD tetrameric interface in Gly-Glu bound N1-N2A receptor structure (6MMP, ref[4]). Residues of K220, L246 in N2A subunits are shown in sticks, with Cα-Cα distances indicated.

**g**, Western blotting analysis on WT and cysteine-substituted N1-N2A receptors. Bands of N2 momomer and N2-N2 homodimer are indicated.

**h,** Cartoon representation of LBD tetrameric interface. Residues of V697, E698, L699 in N1 subunits are shown in sticks, with Cα-Cα distances indicated.

**i**, Assessment of the channel activity on WT and mutant receptors. DTT-induced current amplitude shifts in N1a-N2D (3.56 ± 0.80), N1a$^{E698C}$-N2D (0.65 ± 0.07), N1a$^{E698C}$-N2D$^{L822C}$ (1.33 ± 0.10), N1a$^{C744A,C798A}$-N2D (0.91 ± 0.08), N1a$^{C744A,C798A,V697C}$-N2D (0.73 ± 0.03), N1a$^{C744A,C798A,E698C}$-N2D (0.38 ± 0.08) and N1a$^{C744A,C798A,L699C}$-N2D (0.57 ± 0.13) receptors are shown.

**j,** Evaluation of the relative channel activity in N1a-N2D receptors by MK-801 inhibition kinetics. Representative recording traces and statistics are illustrated in the same manner as in (d). Relative MK-801 $\tau_{on}$ value is 1.00 ± 0.08 for N1a-N2D, 0.62 ± 0.08 for N1a-N2D$^{L822C}$, 0.10 ± 0.01 for N1a$^{E698C}$-N2D and 0.09 ± 0.01 for N1a$^{E698C}$-N2D$^{L822C}$ receptors, n=5-6 oocytes. All values were normalized to the mean value of WT N1a-N2D receptors.

**k.** Assessment of the channel activity on WT and mutant receptors. DTT-induced current amplitude shifts in N1a-N2D (3.56 ± 0.80), N1a$^{E698C}$-N2D (0.65 ± 0.07), N1a$^{E698C}$-N2D$^{L822C}$ (1.33 ± 0.10),

**l**, Western blotting analysis on WT and cysteine-substituted receptors. Bands of N1 momomer, N2D momomer, N1-N1 homodimer and N1-N2D heterodimer are indicated.

**m**, Dose-response curves of Gly and Glu on N1a-N2D (Gly $EC_{50}$ of 0.08 ± 0.01 μM, $n^H$=1.2 and Glu $EC_{50}$ of 0.42 ± 0.04 μM, $n^H$=1.8) and N1a$^{E698C}$-N2D (Gly $EC_{50}$ of 17.49 ± 0.26 μM, $n^H$=2 and Glu $EC_{50}$ of 1.66 ± 0.05 μM, $n^H$=1.7) receptors. n=4 oocyte for each group.

For the statistical analysis, P values are determined by two-tailed unpaired Student's *t*-test for (**d**) and by one-way ANOVA with Tukey's multiple comparison test for (**i-k**) and (***P<0.001; n.s means no significance; error bars, SD).

**Extended Data Fig. 4 | Protein purification, cryo-EM analysis and structure comparison of N1a$^{E698C}$-N2D receptors**

**a-b**, Coomassie blue gel staining and FSEC analysis of purified proteins of N1a$^{E698C}$-N2D receptors. Bands of N1 and N2D monomer and N1-N1 dimer are indicated.

**c**, Cryo-EM data-processing workflow of N1b-N2D receptor datasets processed by Relion 3.1.1[1]. Representative micrographs, 2D average classes, 3D classes and final density maps are shown.

**d,** Local resolution representation of density map and Fourier shell correlation (FSC) of Gly-Glu bound N1b-N2D receptor. Maps are colored based on the local resolution estimation by ResMap-1.1.4[3]. Masked (blue) and unmasked (green) FSCs of corresponding maps are both shown, with gold standard FSC of 0.143 criteria indicated.

**e,** Structural analysis of top-down viewed tetrameric LBD (left), of side-viewed two N1 (middle) and N2D (right) protomers. Center-of-mass (COM) of each lobe, domain and α-helix E (P670-R673 for N1 and R696-Q699 for N2D) is shown in empty circle. Cα atoms of N1 A652 and N2D A678 in gate are marked in filled circle. The dihedral angles for indicating opening-closure degree of LBD are assessed by connecting the Cα of I403, S688, V735, A715 in N1, and P124, E525, S309, E169 in N2D, respectively. Arrows indicate the conformational changes of Gly-Glu bound N1$^{E698C}$-2D C-C state compared to the Gly-Glu bound WT receptor.

**f,** Conformational comparison of N2-LBDs in Gly-Glu bound N1a-N2D, Gly-Glu bound N1a$^{E698C}$-N2D, Gly-Glu bound N1-N2A (PDB:6MMP, ref[4]), Gly-Glu & GNE-6901 bound N1$^{E698C}$-N2A$^{L794C}$ (PDB:7EOR, ref[5]), Gly-Glu bound WT N1b-N2B (PDB:6WI1, ref[6]), Gly-Glu bound N1b$^{E698C}$-N2B$^{L795C}$ (PDB:6WHT, ref[6]) receptors. Dash lines indicate COM distances of D1-D1 and D2-D2 lobes.

**Extended Data Fig. 5 | Biochemical analysis and Cryo-EM data processing of N1-N2C di-receptors in Gly-Glu bound state**

**a**, Cartoon representation of N1-N2C receptor, with mRuby-Strep Tag Ⅱ and 6×His tag placed at the C terminus of N2C and N1 constructs, respectively.

**b**, Coomassie blue gel staining of purified protein of N1-N2C receptors (left panel). FSEC profiles of Gly-Glu bound and Gly-Glu & PYD-106 co-bound N1-N2C receptors (right panel).

**c**, Cryo-EM data processing flowchart of N1-N2C receptors in complex with Gly-Glu. Representative micrographs, 2D class average images and 3D classification maps are shown. Class 3 of best TMD signal was processed individually to get a map displaying certain signal of TMD. Proportion of particle quantity and NTD top-down view comparison of the asymmetric major, intermediate and symmetric minor classes are shown.

**d**, Map alignment of asymmetric major (represented by class 3), intermediate (class 8) and minor (class 9) classes. The rotation angle of one NTD heterodimer is indicated. Middle and right panels show the results of three repeats of 200 ns each unrestrained atomistic simulations on both symmetrical and asymmetrical N1A-N2C structures. The histogram and corresponding kernel density estimation are shown. Middle panel shows the NTD-LBD angles of N2C subunits measured throughout the simulations. Right panel shows RMSD of the least square fitting to the C-alpha atoms, and the first 50ns of simulations in each repeat were excluded for RMSD calculation to allow models to fully relax.

**Extended Data Fig. 6 | N-linked glycosylation analysis by mass spectrometry**

**a**, Sequence alignment of the *Rat norvegicus* N1a, N2A and N2C subunits, highlighted the sites (in purple) detected with N-glycosylation modifications by mass spectrometry.

**b**, Glycans signal (in purple) on EM density maps of N1-N2C (major class) and N1-N2A-N2C receptors are marked. Residue N2C$^{N585}$ located on the intracellular loop between TM1 and TM2 helices, was detected with N-glycosylation modifications, but not present on the map.

**c**, Statistical chart of site-specified N-glycosylation in N1-N2C di-receptors and N1-N2A-N2C tri-receptors by mass spectrometry. The bar plot summarizes the total count of the different N-glycan compositions, and the pie charts summarize the count distribution of different N-glycosylation types for each N-glycosylated site.

**Extended Data Fig. 7 | Cryo-EM data processing of N1-N2C di-receptors in PYD-106 bound state**

Cryo-EM data processing flowchart of N1-N2C receptors in complex with Gly-Glu & PYD-106. Representative micrographs, 2D class average images and 3D classification maps are shown. To push the resolution of ECD, focused refinement was conducted on the PYD-106 bound N1-N2C receptor, with TMD masked out. For the same purpose, one class of best TMD signal was also processed individually to get a map displaying certain signal of TMD. Local resolution estimation and FSC are shown in the same manner as Extended Data Fig.1.

**Extended Data Fig. 8 | Purification, biochemical and cryo-EM analysis of the N1-N2A-N2C tri-receptor**

**a**, Cartoon representation of N1-N2A-N2C tri-receptor, with GFP-6×His tag and GFP-Strep Tag placed at the C terminus of N2A and N2C constructs, respectively.

**b**, Pipeline of two-step affinity chromatography shows that elution of Strep resin was further purified by His resin. Schematic indicates the putative receptor types existed during purification process. Subunit composition at each step was verified by western blotting analysis and the presence of both N2A and N2C subunits was confirmed in the sample after Strep and His affinity purification successively. FSEC profile and Coomassie blue gel staining for the purified tri-receptor protein are shown.

**c**, Flowchart of cryo-EM data processing for Gly-Glu bound N1-N2A-N2C tri-receptor. The initial model was generated *de novo* from the selected 2D particles in Relion 3.1.1[1]. One distinctive 3D class (occupied 11.4% particles) showed N1-N2C di-receptor (major class) liked asymmetric features, which was not considered for final 3D refinement. Local resolution estimation and FSC are shown at the bottom.

**Extended Data Fig. 9 | Cryo-EM density maps and structural comparisons of N1-N2C and N1-N2A-N2C receptors**

**a**, **b**, Electron density of ligands at the LBD clamshells are shown in red mesh (**a**). Representative local densities of N2 subunits in Gly-Glu bound (major class) or PYD-106 bound N1-N2C di-receptors, and N1-N2A-N2C tri-receptor structures (**b**). The models are shown as cartoons and residues are shown as sticks, with N1 coloured in grey, N2A in orange, N2C (chain B) in grey blue and N2C (chain D) in light blue. Agonists Gly and Glu are shown in red sticks.

**c**, Structural comparisons of NTDs and NTD heterodimers within the major class of Gly-Glu bound N1-N2C di-receptor. The r.m.s.d for NTD alignment of N2C (chain B *vs* chain D) and N1 (chain A *vs* chain C) are indicated. NTD heterodimers were superimposed using the R1 lobes of N1 with the rotation angles of R2 lobes indicated.

**d**, Structural comparisons of LBD intra-dimer and inter-dimer with the N1-LBDs aligned within the N1-N2C di- (major class, top panel) or N1-N2A-N2C tri-receptors (bottom panel). Rotation angles of N2-LBDs (from N2C to N2A) in the tri-receptor are indicated. Overall, LBDs exhibit pseudo-symmetry in N1-N2C di-receptor and asymmetry in N1-N2A-N2C tri-receptors

**Extended Data Fig. 10 | Molecular mechanism of PYD-106 selectivity on N1-N2C di-receptors**

**a**, Structural formula of PYD-106 with the only chiral carbon atom in the molecule marked with a red asterisk and the fits of (*R*)- and (*S*)-PYD-106 into the EM map which is shown in mesh. Red arrows indicate the unfavorable fitting for (*S*)-PYD-106.

**b**, Sequence alignment of *Rat norvegicus* N2A, N2B, N2C and N2D subunits. Red boxes indicate the homologous residues at the bottom of the R2 lobe and the top of the D1 lobe, which directly interact with PYD-106 in Ligplot$^+$ in N2C (chain B). Three key residues that form hydrogen bonds with PYD-106 in N2C (R194, D220 and S472) and their homologous residues are highlighted in red.

**c**, Comparison of the NTD-LBD interfaces of N2 subunits in N1-N2C, N1-N2A (PDB:6MMP, ref[4]), N1-N2B (PDB:7EU8, ref[7]) and N1-N2D di-receptors. The hydrophobicity feature of NTD-LBD interface is highlighted with residues at R2 and D1 lobes.

**Supplementary Video 1 | Mechanism of asymmetric allosteric modulation of PYD-106 on N1-N2C di-receptor.**

This movie displays the conformational change of asymmetric N1-N2C di-heteromeric receptor from Gly-Glu to Gly-Glu & PYD-106 bound states, aligned with entire extracellular domains. N1 and N2C subunits are colored in grey and blue, respectively. Upon PYD-106 binding to the N2C of chain B, the relative rotation between NTD and LBD are illustrated.

**Supplementary Video 2 | Molecular organization and architecture of N1-N2A-N2C tri-receptors.**

The video shows the top-down views of asymmetric N1-N2C and N1-N2A (PDB: 6MMP) di-receptors, with N2C (chain B, D) and N2A (chain B, D) colored in blue and orange, respectively. For N1-N2A-N2C tri-receptors, the chain B of N2A and chain D of N2C are preciously integrated into the tri-heteromeric receptor assemble.

**Extended Data Table. 1 | Cryo-EM data collection, refinement and validation statistics for N1-N2D receptors.**

| | N1a-N2D (Gly-Glu) | N1a-N2D (Gly-CPP) | N1a$^{E698C}$-N2D (C-C) | N1a$^{E698C}$-N2D (Non C-C) | N1b-N2D (Gly-Glu) |
|---|---|---|---|---|---|
| **Data collection** | | | | | |
| Magnification | 22500 | 81000 | 81000 | 81000 | 105000 |
| Voltage (kV) | 300 | 300 | 300 | 300 | 300 |
| Electron exposure ($e^{-}/Å^2$) | 60 | 60 | 60 | 60 | 60 |
| Defocus range (μm) | -1.5 ~ -2.5 | -1.5 ~ -2.5 | -1.5 ~ -2.5 | -1.5 ~ -2.5 | -2.0 ~ -2.5 |
| Pixel size (Å) | 1.067 | 1.071 | 1.071 | 1.071 | 0.803 |
| Symmetry imposed | C2 | C2 | C2 | C2 | C2 |
| Initial particles | 505,204 | 535,387 | 470,935 | 470,935 | 678,197 |
| Final particles | 232,194 | 142,841 | 91,328 | 143,824 | 98,449 |
| Map resolution (Å) | 4.0 | 3.7 | 6.4 | 4.3 | 5.1 |
| FSC threshold | 0.143 | 0.143 | 0.143 | 0.143 | 0.143 |
| Map resolution range (Å) | 3.88-10.78 | 3.18-8.70 | 4.85-12.34 | 3.45-11.37 | 3.85-10.24 |
| **Refinement** | | | | | |
| Initial model (PDB code) | 6WI1 | 6WI1 | 6WI1 | 6WI1 | 6WI1 |
| Map sharpening *B* factor ($Å^2$) | -220 | -150 | -220 | -250 | -230 |
| Model composition | | | | | |
| Non-hydrogen atoms | 20582 | 22422 | 17922 | 20018 | 19014 |
| Protein residues | 2804 | 3020 | 2538 | 2674 | 2518 |
| Ligands | 24 | 20 | 12 | 48 | 12 |
| *B* factors ($Å^2$) | | | | | |
| Protein | 35.37 | 68.87 | 416.14 | 109.95 | 234.37 |
| Ligand | 20 | 112.20 | 20 | 20 | 20 |
| R.m.s. deviations | | | | | |
| Bond lengths (Å) | 0.002 | 0.003 | 0.003 | 0.017 | 0.003 |
| Bond angles (°) | 0.587 | 0.611 | 0.586 | 0.632 | 0.697 |
| **Validation** | | | | | |
| MolProbity score | 1.80 | 1.98 | 2.13 | 1.85 | 2.21 |
| Clashscore | 6.47 | 8.68 | 12.95 | 8.69 | 13.69 |
| Poor rotamers (%) | 0.42 | 0 | 0 | 0 | 0 |
| Ramachandran plot | | | | | |
| Favored (%) | 93.02 | 91.29 | 91.3 | 94.44 | 89.36 |
| Allowed (%) | 6.84 | 8.71 | 8.46 | 5.56 | 10.56 |
| Disallowed (%) | 0.15 | 0 | 0.24 | 0 | 0.08 |

**Extended Data Table. 2 | Cryo-EM data collection, refinement and validation statistics for N2C-containing receptors.**

| | N1a-N2C (major class, Gly-Glu) | N1a-N2C (minor class, Gly-Glu) | N1a-N2C (Gly-Glu & PYD-106) | N1a-N2A-N2C (Gly-Glu) |
|---|---|---|---|---|
| **Data collection** | | | | |
| Magnification | 81,000 | 81,000 | 81,000 | 81,000 |
| Voltage (kV) | 300 | 300 | 300 | 300 |
| Electron exposure (e$^{-}$/Å$^{2}$) | 60 | 60 | 60 | 60 |
| Defocus range (μm) | -1.2 ~ -2.0 | -1.2 ~ -2.0 | -1.2 ~ -2.0 | -1.2 ~ -2.0 |
| Pixel size (Å) | 1.071 | 1.071 | 1.071 | 1.071 |
| Symmetry imposed | C1 | C2 | C1 | C1 |
| Initial particles | 1,188,229 | 1,188,229 | 2,094,482 | 891,914 |
| Final particles | 245,730 | 15,479 | 601,826 | 278,030 |
| Map resolution (Å) | 3.61 | 4.31 | 3.03 | 3.53 |
| FSC threshold | 0.143 | 0.143 | 0.143 | 0.143 |
| Map resolution range (Å) | 3.41-6.21 | | 2.89-4.71 | 3.08-7.20 |
| **Refinement** | | | | |
| Initial model (PDB code) | 6WI1 | 3WI1 | 6WI1 | 6MMT |
| Map sharpening *B* factor (Å$^{2}$) | -125 | -134 | -50 | -125 |
| Model composition | | | | |
| Non-hydrogen atoms | 21222 | 20901 | 21289 | 24138 |
| Protein residues | 2620 | 2608 | 2620 | 2980 |
| Ligands | 48 | 32 | 53 | 45 |
| *B* factors (Å$^{2}$) | | | | |
| Protein | 38.18 | 278.15 | 90.65 | 135.75 |
| Ligand | 81.14 | 333.80 | 140.15 | 102.90 |
| R.m.s. deviations | | | | |
| Bond lengths (Å) | 0.011 | 0.003 | 0.007 | 0.003 |
| Bond angles (°) | 1.113 | 0.703 | 0.808 | 0.672 |
| **Validation** | | | | |
| MolProbity score | 1.94 | 2.21 | 1.79 | 1.96 |
| Clashscore | 10.43 | 19.91 | 7.48 | 8.80 |
| Poor rotamers (%) | 0.18 | 0.00 | 0.13 | 1.20 |
| Ramachandran plot | | | | |
| Favored (%) | 93.92 | 93.71 | 94.42 | 93.59 |
| Allowed (%) | 5.88 | 6.02 | 5.35 | 6.00 |
| Disallowed (%) | 0.19 | 0.27 | 0.23 | 0.41 |
