## Supplemental figures and tables for "Distinct structure and gating mechanism in diverse NMDA receptors with GluN2C and GluN2D subunits"

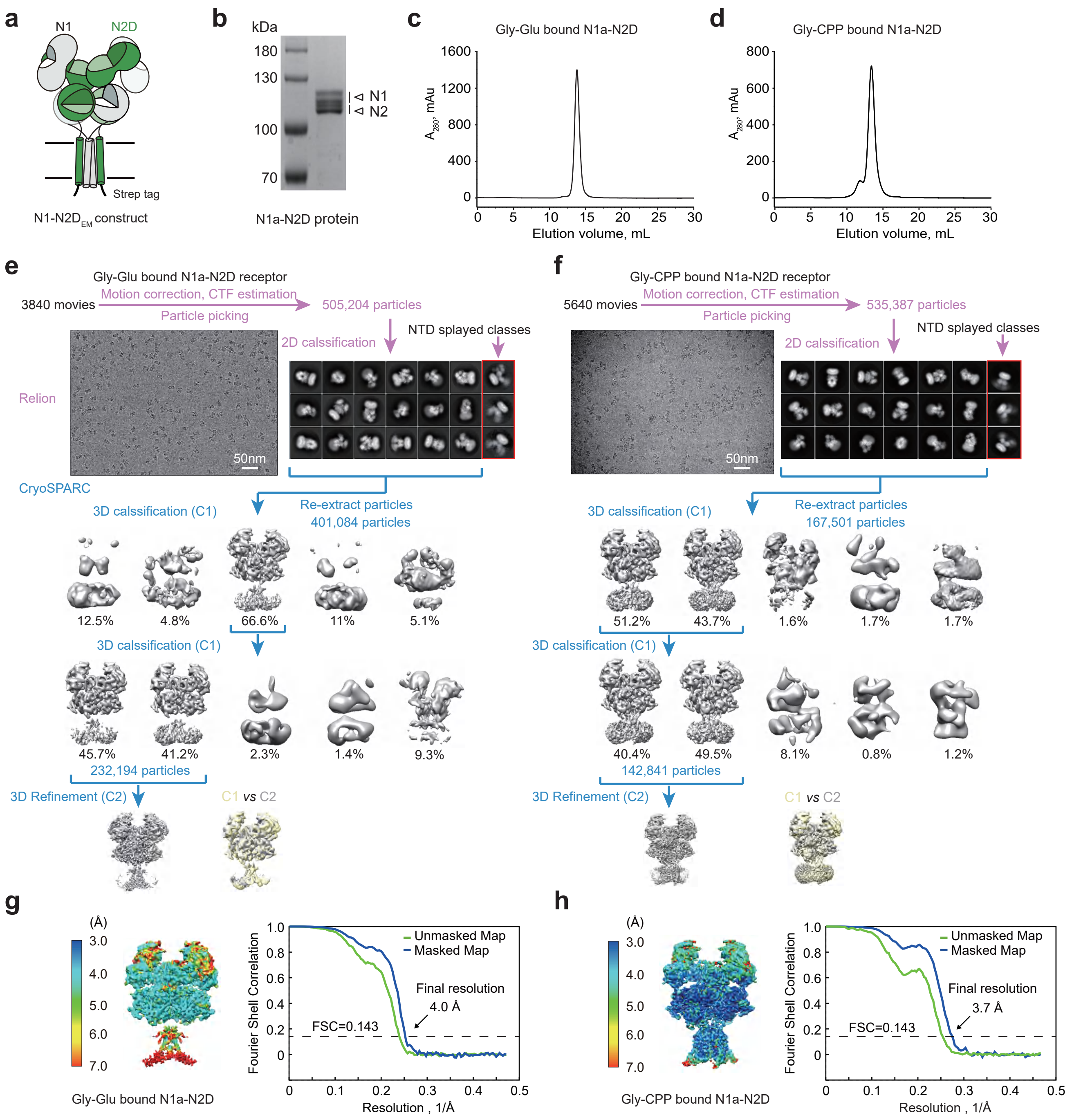


**Extended Data Figure. 1**

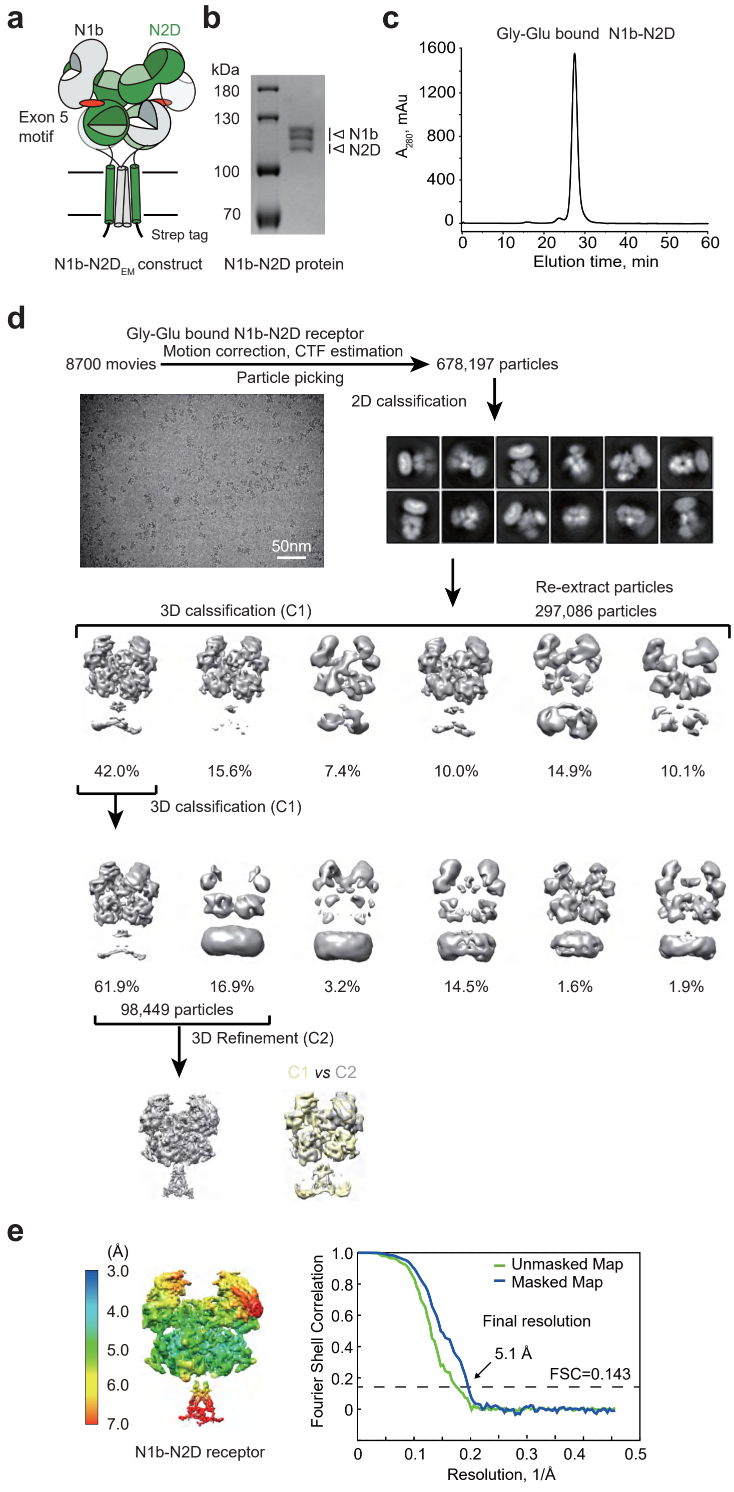


**Extended Data Figure. 2**

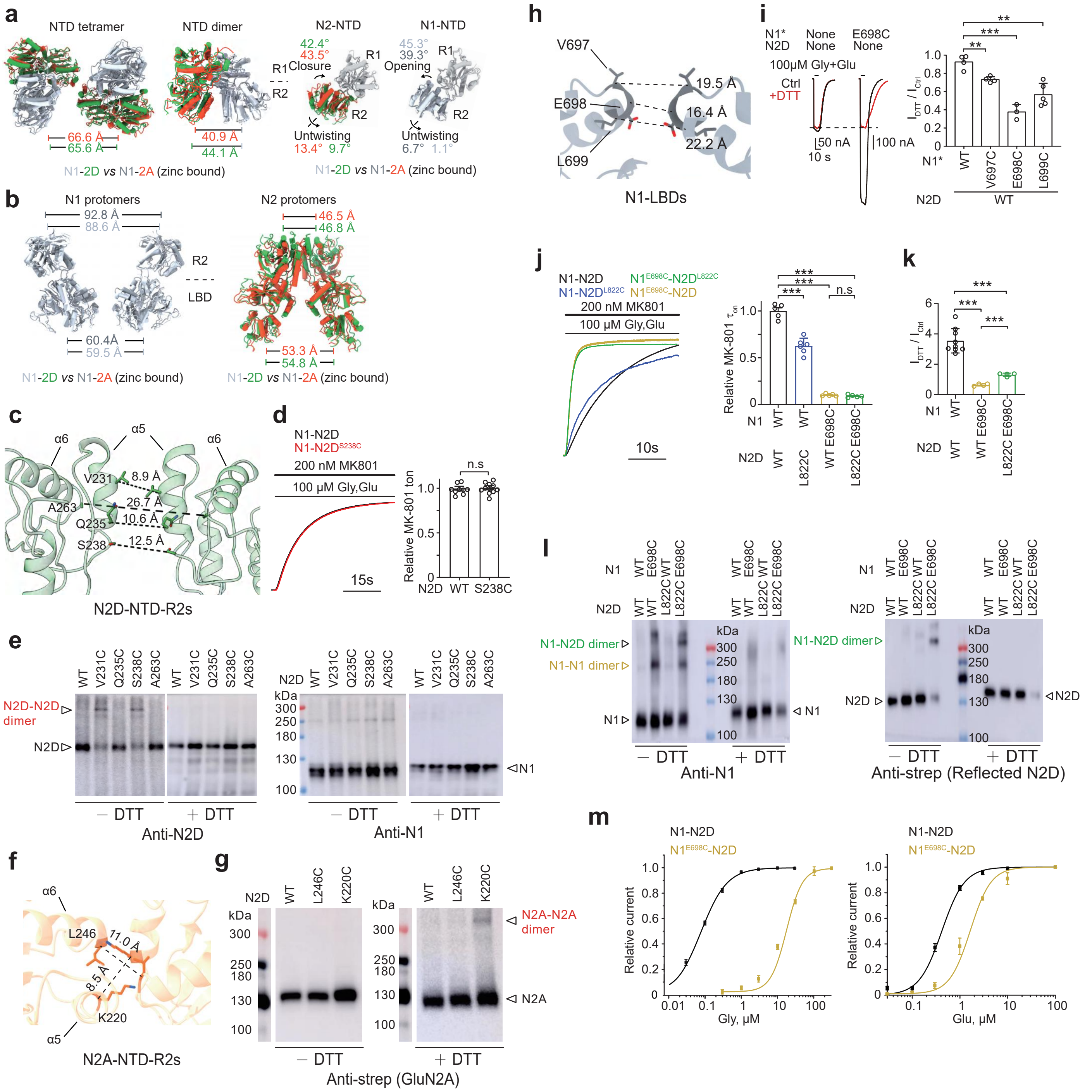


**Extended Data Figure. 3**

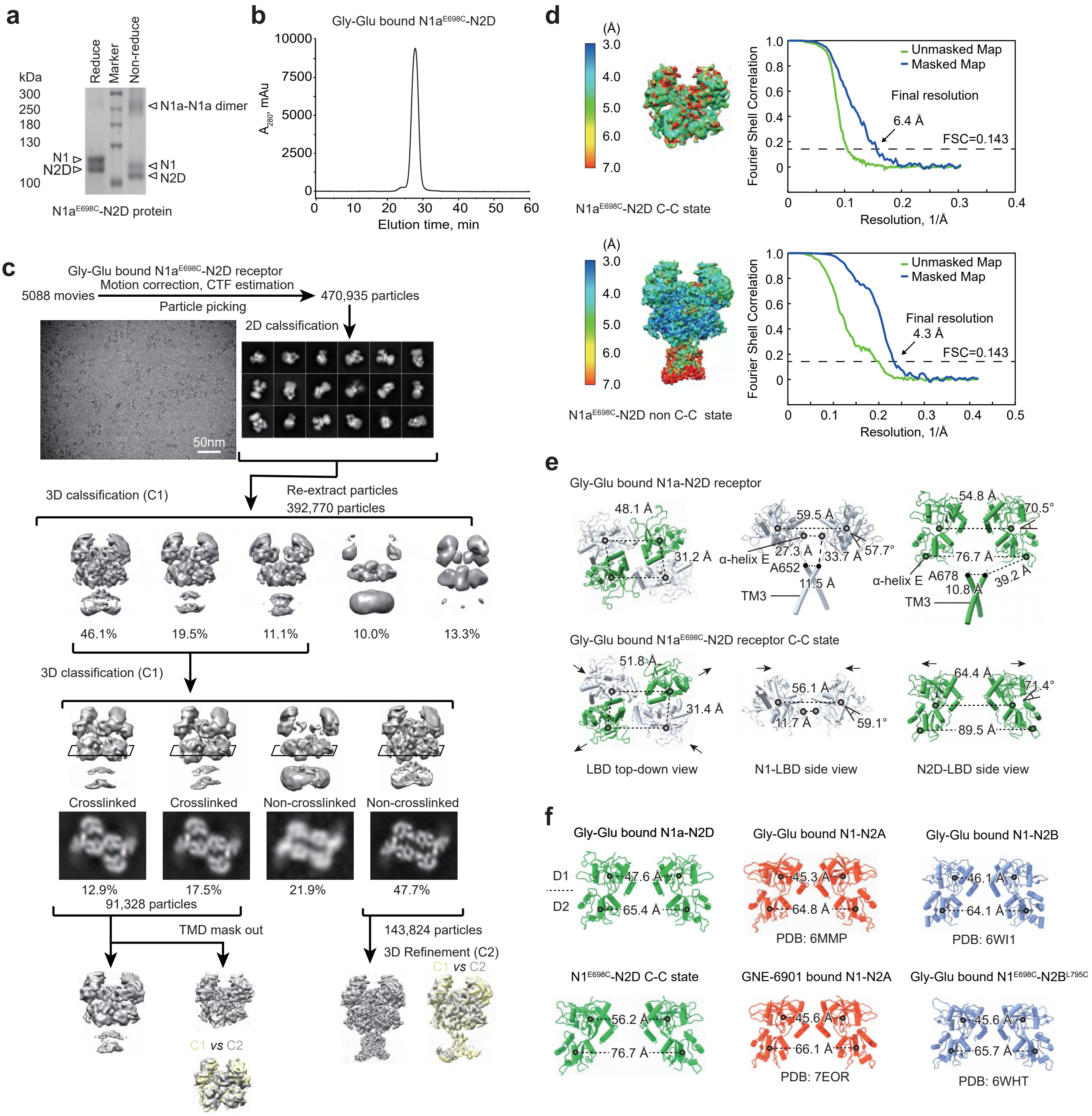


**Extended Data Figure. 4**

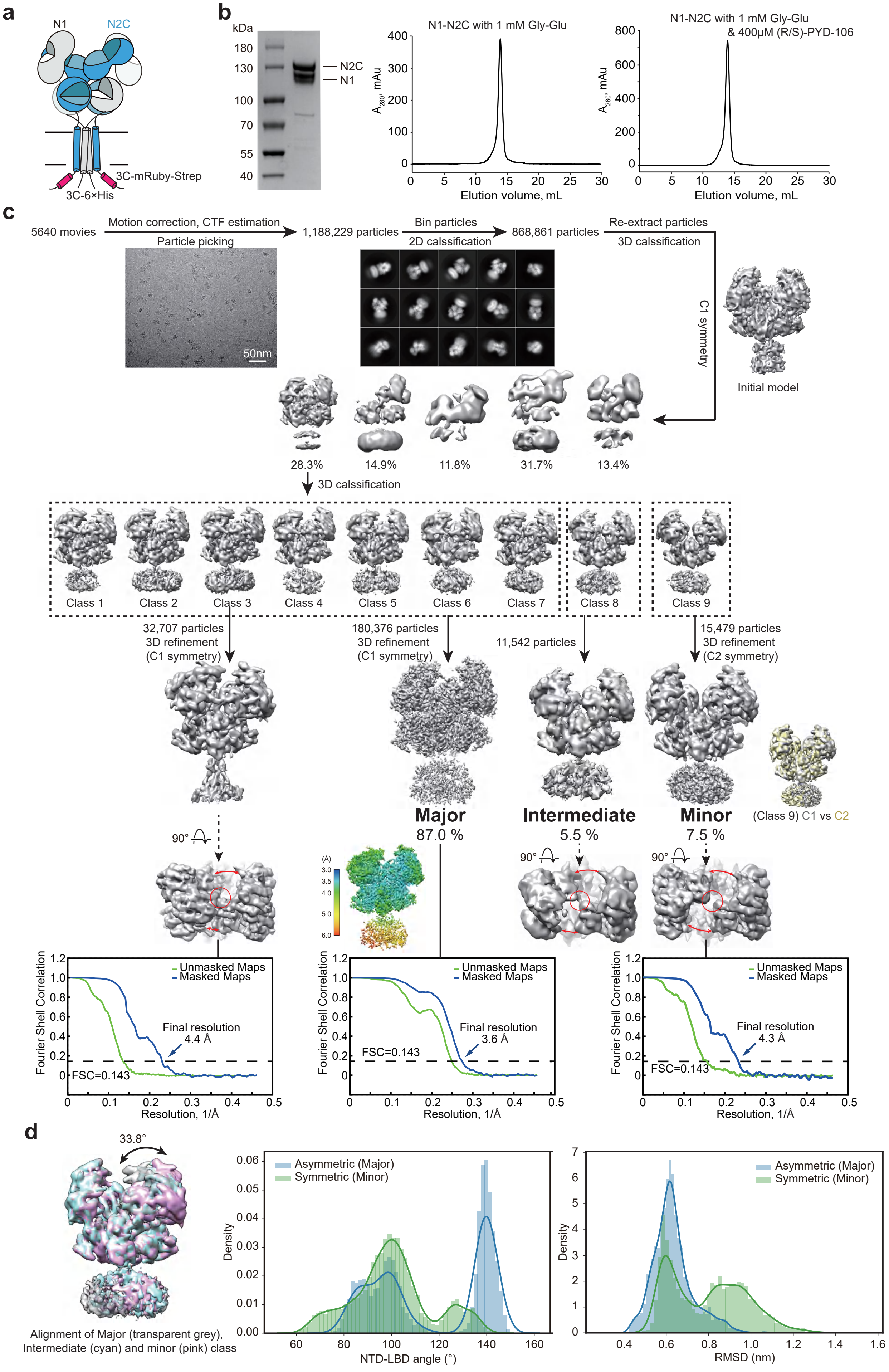


**Extended Data Figure. 5**

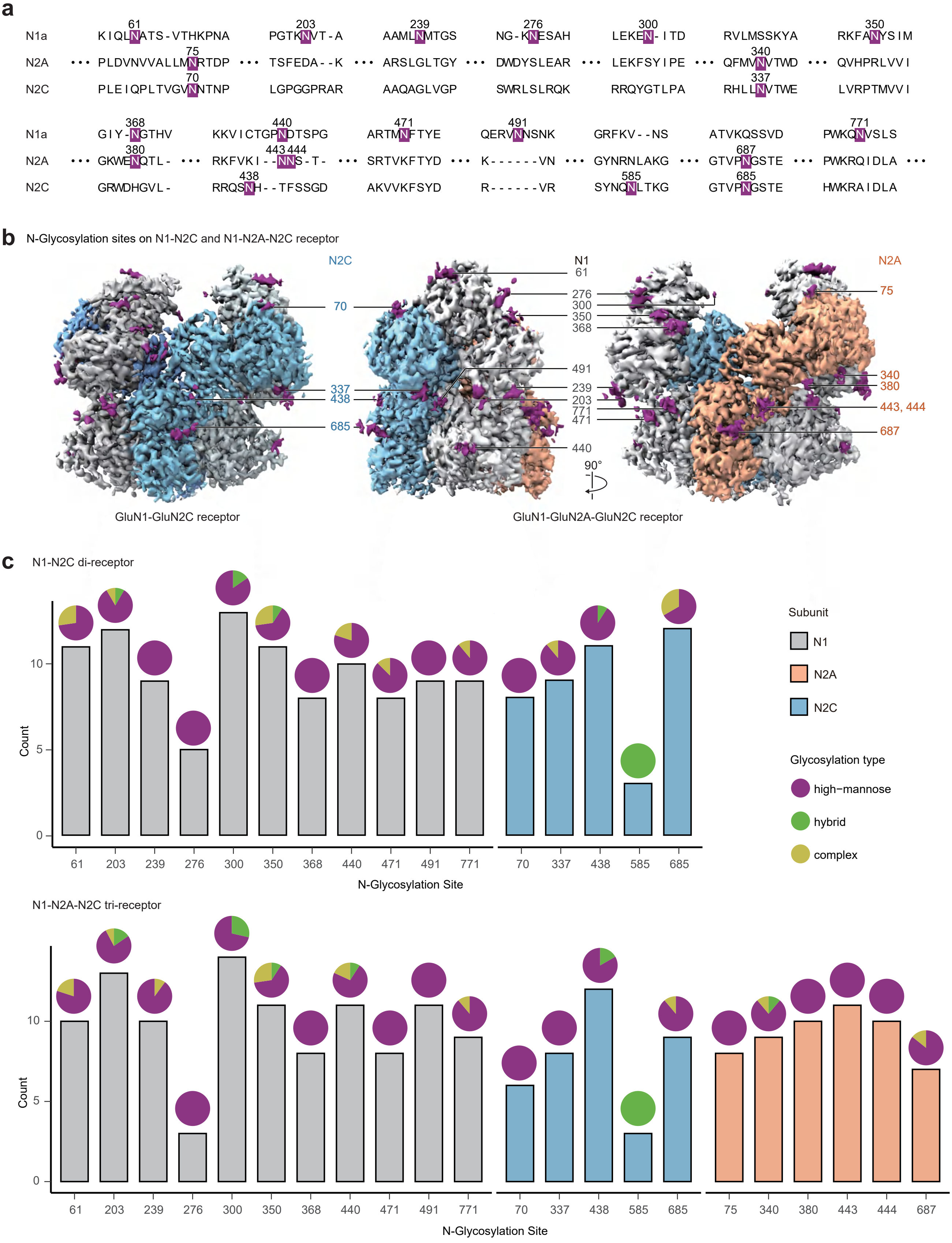


**Extended Data Figure. 6**

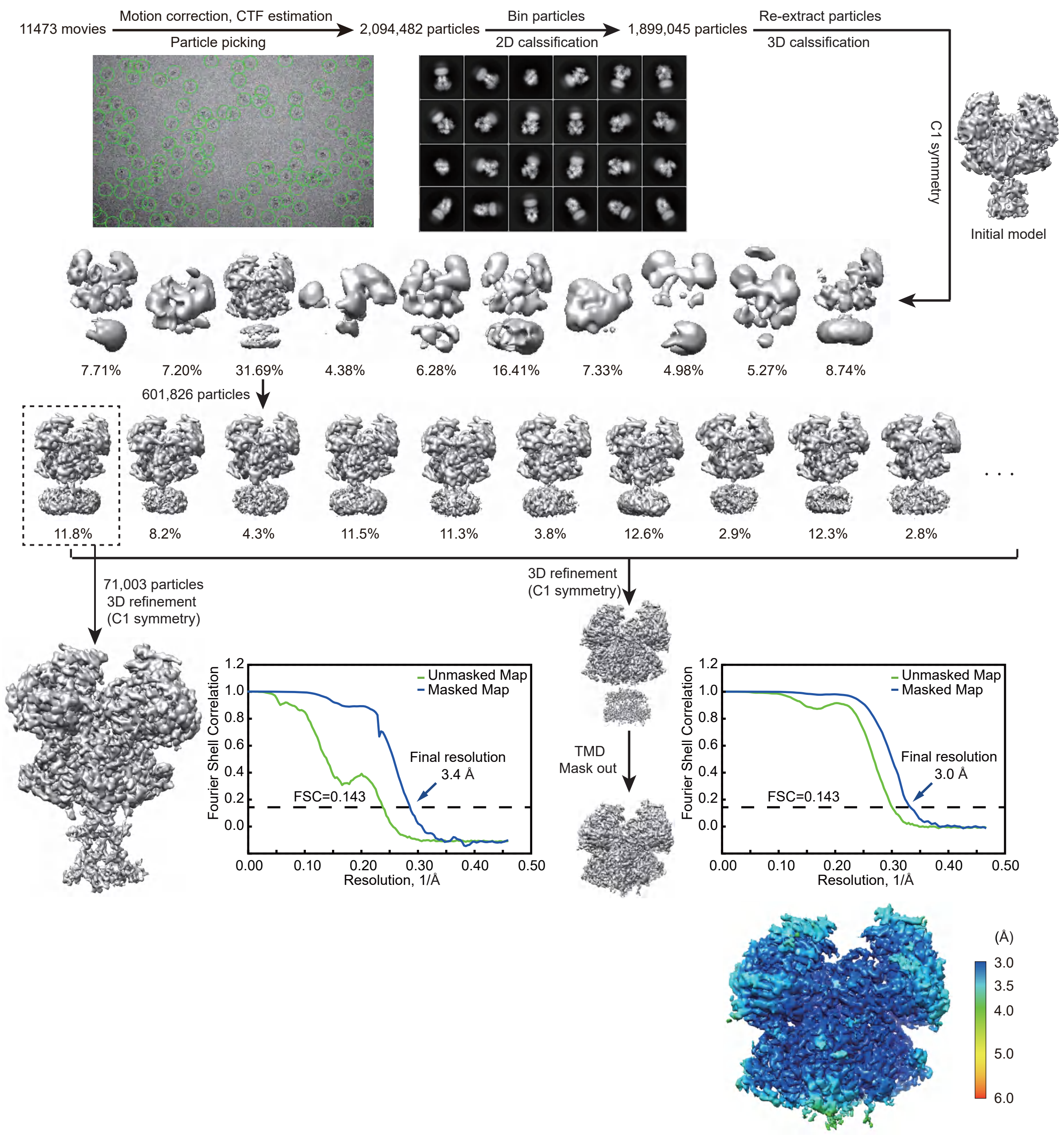


**Extended Data Figure. 7**

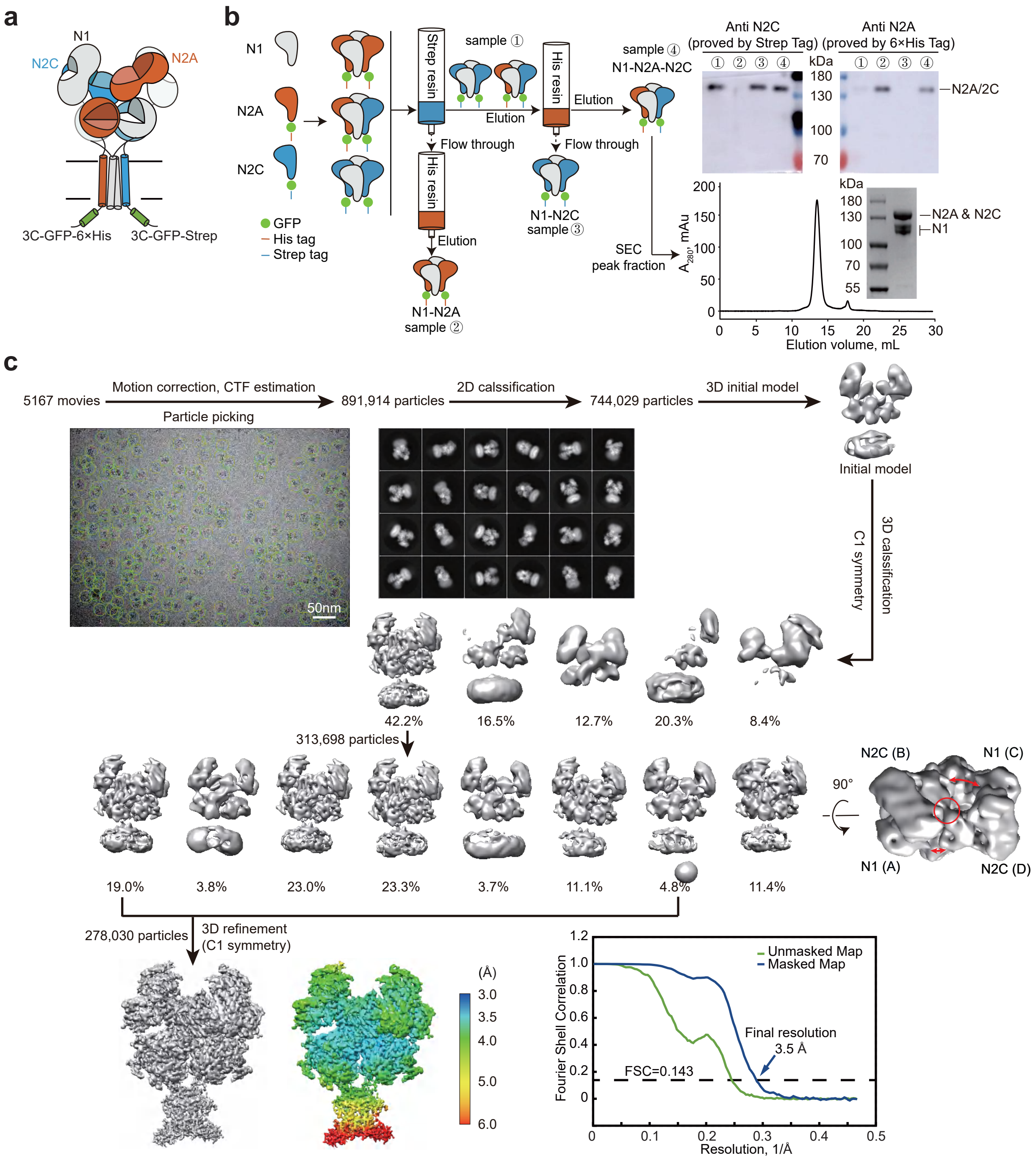


**Extended Data Figure. 8**

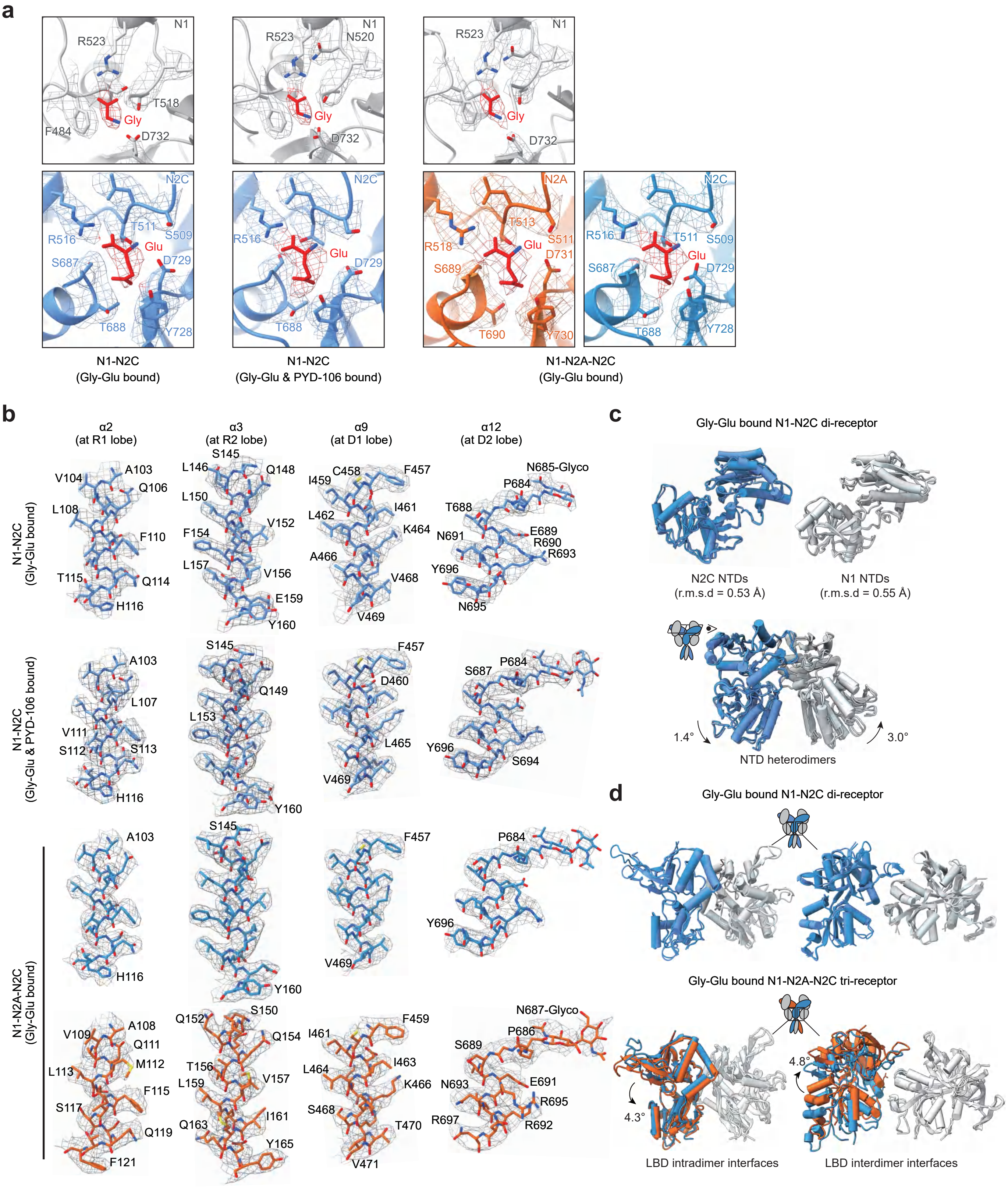


#### Extended Data Figure. 9

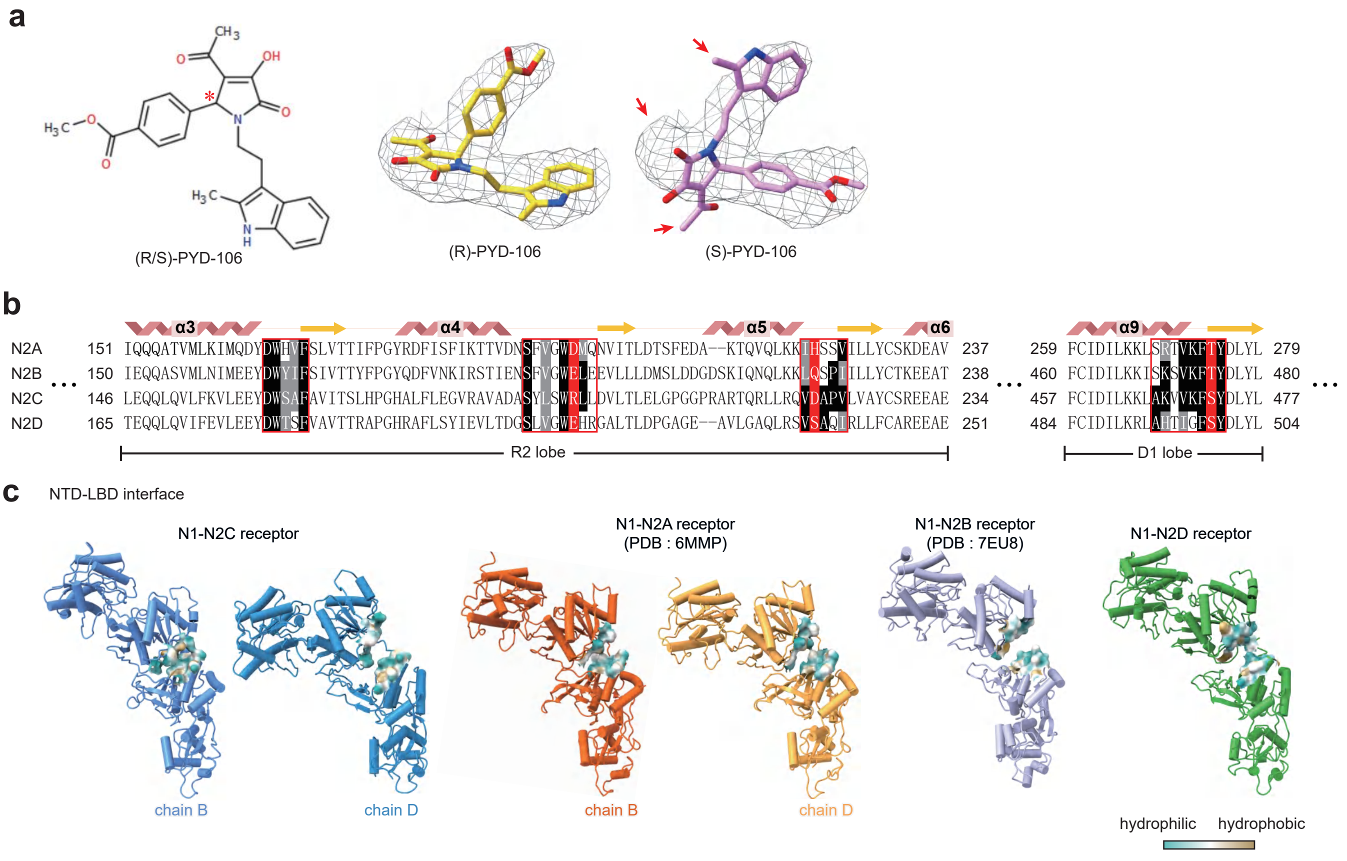


### Extended Data Figure. 10
